# IZUMO1 is a sperm fusogen

**DOI:** 10.1101/2022.02.01.478669

**Authors:** Nicolas G. Brukman, Kohdai P. Nakajima, Clari Valansi, Kateryna Flyak, Xiaohui Li, Tetsuya Higashiyama, Benjamin Podbilewicz

## Abstract

Mammalian sperm-egg adhesion depends on the trans-interaction between the sperm-specific type I glycoprotein IZUMO1 and its oocyte-specific GPI-anchored receptor JUNO. However, the mechanisms and proteins (fusogens) which mediate the following step of gamete fusion remain unknown. Using live imaging and content mixing assays in a heterologous system and structure-guided mutagenesis, we unveil an unexpected function for IZUMO1 in cell-to-cell fusion. We show that IZUMO1 alone is sufficient to induce fusion, and that this ability is retained in a mutant unable to bind JUNO. On the other hand, a triple mutation in exposed aromatic residues prevents this fusogenic activity without impairing JUNO interaction. Our findings suggest a second, crucial function for IZUMO1 as a unilateral mouse gamete fusogen.

**Highlights:**

- IZUMO1 expression in somatic cells in culture induces cell-to-cell fusion
- The fusogenic activity of IZUMO1 is unilateral
- Cell fusion is independent of the binding of IZUMO1 to JUNO
- IZUMO1-mediated cell merger depends on its transmembrane domain, and three solvent-exposed aromatic residues

**Graphical abstract:** 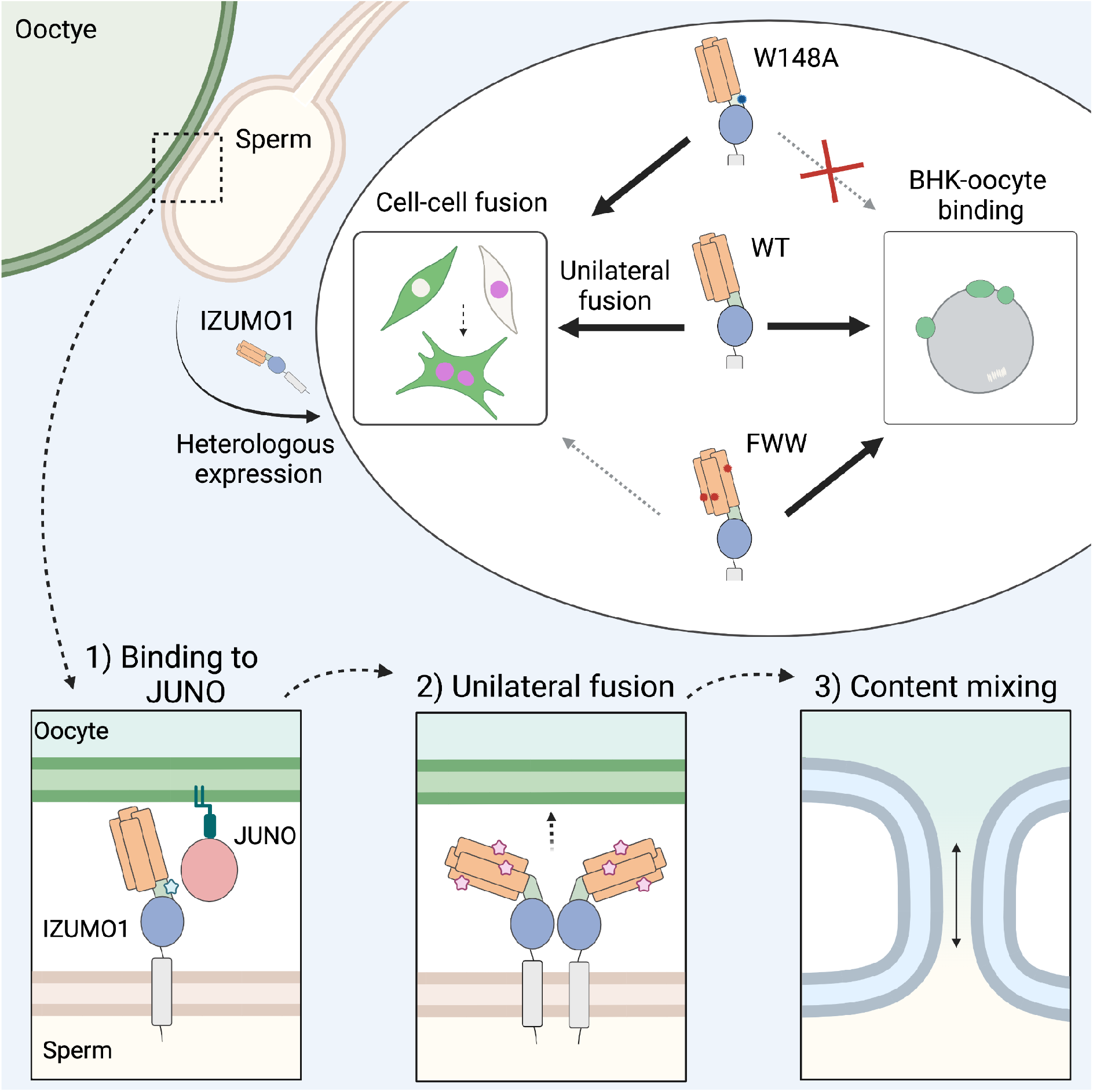

## Introduction

The final steps of mammalian egg-sperm fusion remain a mechanistic enigma. Cell-to-cell fusion requires the action of specialized proteins, named fusogens, to overcome the energetic barriers that arise when two plasma membranes come into close proximity (Chernomordik and Kozlov, 2003). By definition, fusogens are both necessary in their system of origin (*in situ*) and sufficient to induce membrane merging in otherwise non-fusing heterologous systems (Segev et al., 2018). While a mystery in mammals, gamete fusion in flowering plants and protists is mediated by the fusogen GENERATIVE CELL-SPECIFIC 1/HAPLESS 2 (GCS1/HAP2). The essentiality of GCS1/HAP2 in gamete fusion was first demonstrated in *Arabidopsis thaliana*, being sperm-expressed and necessary for sperm-egg and sperm-central cell fusion (von Besser et al., 2006; Johnson et al., 2004; Mori et al., 2006). GCS1/HAP2 is also essential to fuse gametes in the malaria parasite *Plasmodium*, in the slime mold *Dictyostelium* and in the algae *Chlamydomonas* (Hirai et al., 2008; Liu et al., 2008; Okamoto et al., 2016). We have subsequently demonstrated that the expression of *Arabidopsis thaliana* GCS1/HAP2 is sufficient to fuse mammalian cells in culture (Valansi et al., 2017), thereby characterizing this protein as a bona fide fusogen. Remarkably, structural analysis of GCS1/HAP2 from different species showed a shared tertiary and quaternary organization with class II viral glycoproteins (such as those from dengue, rubella and zika viruses) (Fédry et al., 2017; Pinello et al., 2017; Valansi et al., 2017) and with Fusion Family (FF) proteins from nematodes and other organisms (Avinoam et al., 2011; Mohler et al., 2002; Pérez-Vargas et al., 2014; Sapir et al., 2007). This protein superfamily, termed Fusexins (Valansi et al., 2017), are widely distributed in multiple phyla, but to date no members have been identified in vertebrates (Brukman et al., 2022; Vance and Lee, 2020).

Unlike gamete fusogens, other mammalian somatic fusogens are known and well-characterized. For example, fusion of myoblasts to form and maintain muscle fibers requires the coordinate action of Myomaker (TMEM8c) and Myomerger (Myomixer/Minion/Gm7325) (Bi et al., 2017; Millay et al., 2013; Quinn et al., 2017; Zhang et al., 2017). Their expression in fibroblasts drives cell-to-cell fusion: Myomerger can work unilaterally from either one of the merging membranes, while Myomaker is required on both fusing cells (bilateral mechanism) (Leikina et al., 2018). During placenta formation, extensive trophoblast fusion is mediated by syncytins. Remarkably, syncytins are evolutionarily related to retroviral Class I glycoproteins (Lavialle et al., 2013). *Syncytin-A* and *-B* mutations in mice results in fusion defects during the formation of the syncytiotrophoblast (Dupressoir et al., 2009, 2011); while human Syncytin-1 or -2 expression is sufficient to induce heterologous fusion of cells in culture (Blond et al., 2000; Esnault et al., 2008). Interestingly, the fusogenic activity of Syncytin-1 and -2 was highly increased when the respective receptor, ASCT2 or MFSD2, was present (Blond et al., 2000; Esnault et al., 2008). A tight binding step to a receptor in the target cell preceding membrane fusion is a common mechanism among viral fusogens, with binding and fusion often mediated by different domains or subunits within the fusogen (Podbilewicz, 2014; White et al., 2008).

A number of events preceding gamete fusion take place during fertilization. Sperm must undergo a physiological process called capacitation, which includes the exocytosis of the acrosome, a specialized vesicle in the head (Yanagimachi, 1994). This allows the sperm to penetrate a proteinic coating which covers the oocyte, called vitelline envelope or, in mammals, *zona pellucida* (ZP) (Wassarman, 1999; Yanagimachi, 1994). Only after penetration of the ZP, the plasma membranes of both gametes are able to bind to each other and finally fuse to complete fertilization (Bianchi and Wright, 2020). Some proteins expressed in the oocyte or in the sperm were shown to be essential for the last steps of fertilization (Deneke and Pauli, 2021). The oocyte tetraspanins CD9 and CD81 are required for sperm-egg fusion (Kaji et al., 2000; Le Naour et al., 2000; Miyado et al., 2000; Rubinstein et al., 2006) by regulating membrane architecture and compartmentalization (Inoue et al., 2020; Runge et al., 2007). Mutation of any of the sperm-specific proteins TMEM95, SPACA6, FIMP, SOF1 and DCST1/2 leads to male infertility due to defects in gamete fusion (Barbaux et al., 2020; Fujihara et al., 2020; Inoue et al., 2021; Lamas-Toranzo et al., 2020; Lorenzetti et al., 2014; Noda et al., 2020, 2022). While all these genes are essential for late stages in fertilization, and loss-of-function mutations of any of them prevent gamete fusion, it is not clear what specific step in the process is being directly affected (Brukman et al., 2019). The only known pair of trans-interacting proteins are IZUMO1, in the sperm, and JUNO/IZUMO1R, from the oocyte. IZUMO1 was the first characterized sperm protein whose deletion was demonstrated to block gamete fusion in mouse (Inoue et al., 2005), while JUNO was subsequently identified as the IZUMO1 receptor in the oocyte (Bianchi et al., 2014). The IZUMO1-JUNO interaction is required for efficient sperm-egg binding (Matsumura et al., 2021) in a species-specific manner (Bianchi and Wright, 2015). Nevertheless, a previous study failed to detect fusion between mixed Human Embryonic Kidney HEK293T cells expressing JUNO or IZUMO1 (Bianchi et al., 2014). Here, however, we present evidence that the expression of mouse IZUMO1 is sufficient to mediate cell-to-cell fusion independently of its binding to JUNO, thus underpinning an additional role for IZUMO1 in membrane merger during fertilization.

## Results

### IZUMO1 is sufficient to induce syncytia formation in somatic cells

There are two main attributes that define a membrane fusogen: First, the molecule has to be necessary for membrane fusion so when it is absent or mutated fusion fails to occur; and second, the protein must be sufficient to merge cells that normally do not fuse. Generally non-fusing cells in culture, such as Baby Hamster Kidney (BHK) cells, have been widely used to test the fusogenic ability of proteins and illustrate fusogen sufficiency (Figure S1A, (Avinoam et al., 2011; Bitto et al., 2016; Jeetendra et al., 2002; Valansi et al., 2017; Waning et al., 2004; White et al., 1981)). To look for gamete fusogen(s) in mammals, we aimed to evaluate candidate gamete-specific transmembrane proteins using this system, with mouse egg JUNO and sperm IZUMO1 as negative controls to account for trans-interacting proteins involved in binding but not fusion. To our surprise, when we expressed IZUMO1, but not JUNO, we observed syncytia formation at levels similar to those in a positive control using GCS1/HAP2, a gamete fusogen from *Arabidopsis thaliana* (Figure 1A). We found that multinucleation increased three-fold when IZUMO1 was expressed compared to the negative control containing a membrane-bound myristoylated EGFP (myrGFP) (Figure 1B). For both IZUMO1 and GCS1/HAP2, we found an increase in cells with two to five-nuclei (Table S1). The presence of both IZUMO1 and JUNO, on the surface of BHK cells was confirmed by immunostaining of non-permeabilized samples (Figure S2).

**Figure 1.**
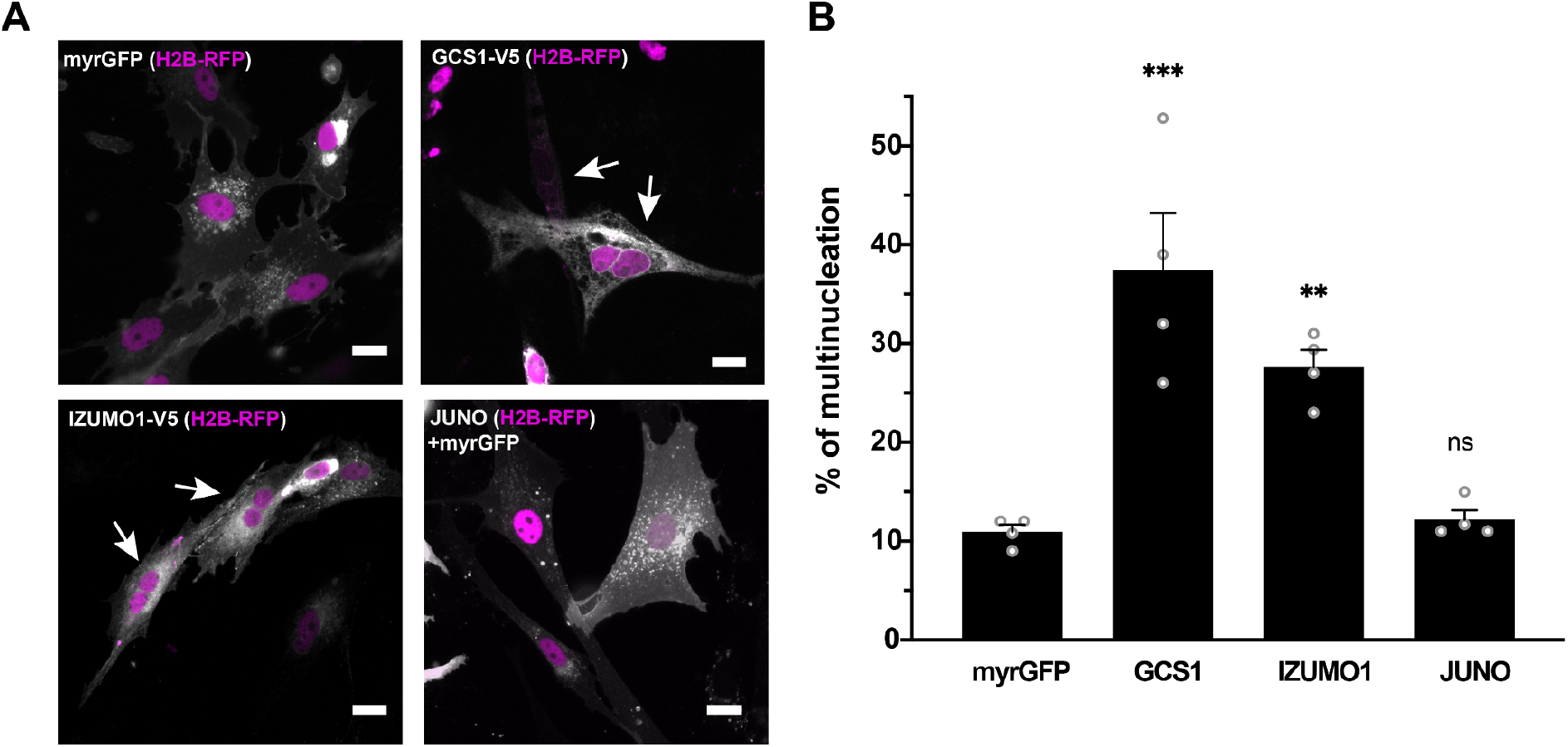
IZUMO1 induces multinucleation of BHK cells. **(A)** Cells were transfected with either pCI::myrGFP::H2B-RFP (myristoylated EGFP), pCI::GCS1/HAP-V5::H2B-RFP, pCI::IZUMO1-V5:H2B-RFP or pCI::JUNO::H2B-RFP vectors. JUNO was co-transfected with a plasmid for myrGFP. Immunofluorescence was performed with anti-V5 antibodies for IZUMO1 and GCS1/HAP2, while the myrGFP signal is shown for JUNO and vector control (gray). Arrows show cells with more than one nucleus (magenta). Scale bars, 20 µm. **(B)** The percentage of multinucleation (see Supplementary Information) is presented as individual data and means ± SEM of four independent experiments. The number of cells counted can be found in Table S1. Comparisons were made with one-way ANOVA followed by Dunett’s test against the empty vector. ns = non-significant, ** p < 0.01, *** p < 0.001.

### IZUMO1 is sufficient to fuse cells

Following the observation that IZUMO1 can induce multinucleation, we aimed to test the ability of IZUMO1 and JUNO to mediate cell-to-cell fusion by using content mixing experiments (Avinoam et al., 2011; Valansi et al., 2017). In this assay, two populations of BHK cells expressing either cytosolic EGFP (GFPnes) or nuclear H2B-RFP are co-incubated, and the appearance of multinucleated cells containing both fluorescent markers is registered as an indication of fusion (Figures 2A and S3). Under these conditions, expression of JUNO alone failed to induce content mixing, consistent with the syncytia formation assay. In contrast, IZUMO1 induced a 5.5-fold increase in the content mixing of BHK cells compared to the controls employing the fluorescent empty vectors; while a 4-fold increase was observed in the positive GCS1/HAP2 control (Figures 2B, S3 and S4A). To determine whether IZUMO1 can merge different heterologous cells, we performed content mixing assays using human HEK293T cells and obtained similar results (Figure S5). Thus, the mouse sperm protein IZUMO1 is sufficient to fuse hamster and human cells; whereas the egg receptor of IZUMO1 (JUNO) is unable to fuse heterologous cells (Figure S1B).

**Figure 2.**
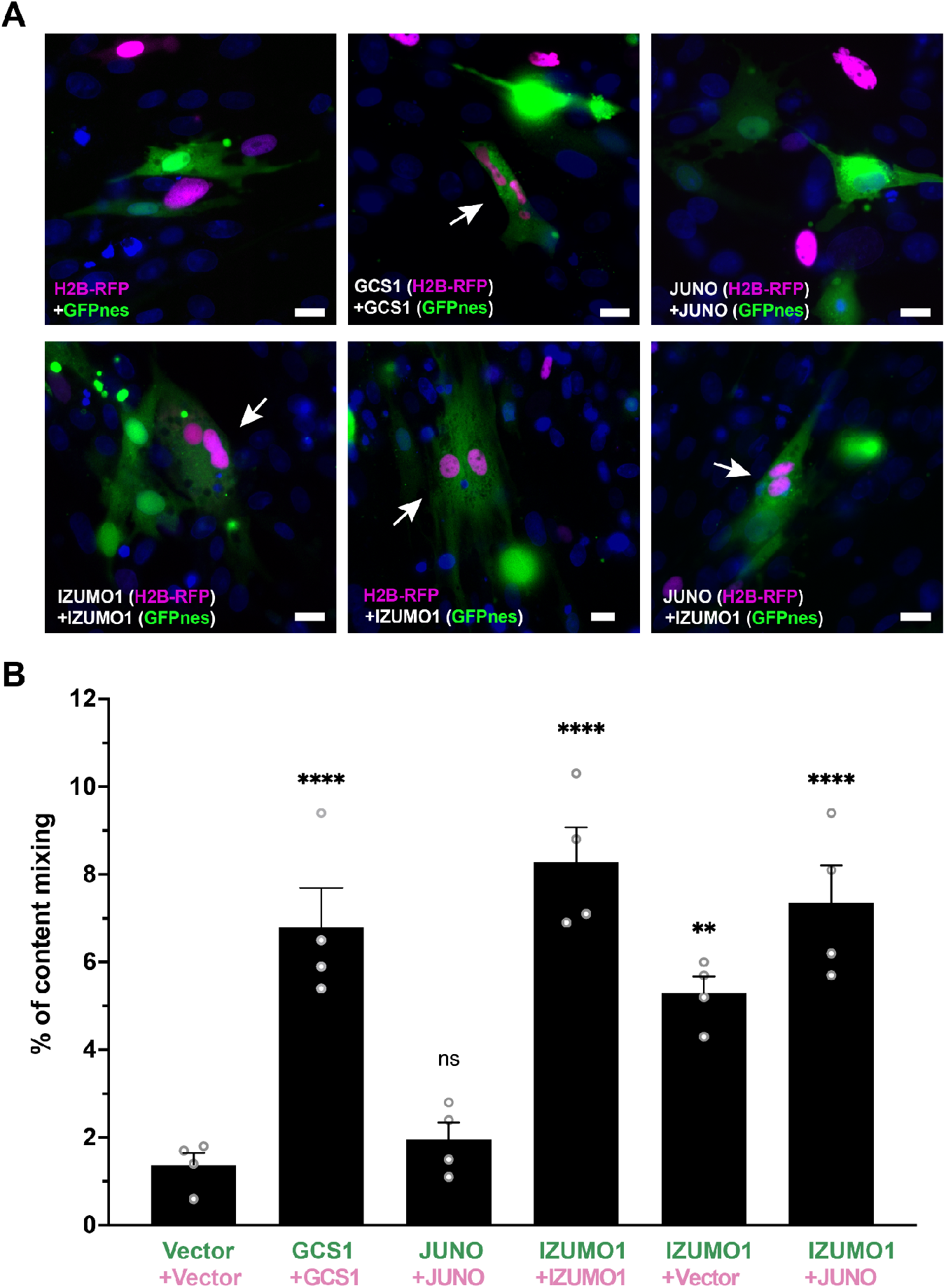
IZUMO1 induces fusion of BHK cells. **(A)** Representative images of mixed cells transfected with pCI::GFPnes or pCI::H2B-RFP empty vectors or containing the coding sequence for the expression of GCS1/HAP2, IZUMO1 and JUNO as indicated. Arrows show content-mixed cells containing both GFPnes (green cytoplasm) and H2B-RFP (red nuclei). DAPI staining is shown in blue. Scale Bars, 20 µm. See also Figure S3. **(B)** Quantification of content-mixing experiments. The percentage of mixing was defined as the ratio between the nuclei in mixed cells (NuM) and the total number of nuclei in mixed cells and fluorescent cells in contact that did not fuse (NuC), as follows: % of mixing = (NuM/(NuM+NuC)) x 100. Bar chart showing individual experiment values (each corresponding to 1000 nuclei) and means ± SEM of four independent experiments. Comparisons by one-way ANOVA followed by Dunett’s test against the empty vectors. ns = non-significant, ** p < 0.01, **** p < 0.0001.

### IZUMO1-mediated fusion uses a viral-like unilateral mechanism

Some fusogens, like the hemagglutinin of influenza virus or the syncytins, can mediate fusion when they are expressed in only one of the fusing membranes while others, like EFF-1 and AFF-1 from nematodes, are required in both fusing membranes acting in a bilateral way (Blond et al., 2000; Esnault et al., 2008; Podbilewicz et al., 2006; White et al., 1982). As IZUMO1 is present only in the sperm we aimed to test whether cells expressing IZUMO1 are able to fuse to cells transfected with an empty vector or expressing JUNO. We found that content mixing was observed when IZUMO1 was present only in one of the two populations of cells that were mixed, implying a unilateral fusion mechanism like in viruses (Figures 2 and S3). Expressing JUNO *in trans* of IZUMO1 appeared to increase the number of fusion events, which may be due to increased association between cells from the two populations as a result of IZUMO1-JUNO interaction. In summary, our results suggest that mouse gamete fusion can be mediated by the viral-like unilateral fusogenic activity of IZUMO1 on the sperm that follows the docking mediated by the IZUMO1-JUNO interaction (Figure S1B).

### Time-lapse imaging reveals the dynamics of IZUMO1-mediated fusion

To independently study the apparent fusogenic activity of IZUMO1, we performed live imaging experiments to track fusion events of BHK cells in real time using an inducible system (Figure S1A). For this purpose, we transfected cells with a plasmid encoding for cytoplasmic RFP (RFPcyto) alone, for the negative control, or together with a mifepristone-inducible system to express our candidate proteins. As a positive control we used EFF-1, a somatic fusogen from *Caenorhabditis elegans* (Avinoam et al., 2011). Vectors for E-cadherin, GEX2 (Gamete EXpressed 2), IZUMO1 and JUNO were similarly prepared and the cells were visualized upon induction. We observed that cell-to-cell fusion occurred following EFF-1 expression (Figure 3A; Movie S1). Cell-to-cell fusion events were quantified 12 h after induction. We found that EFF-1 expression significantly increased fusion levels (Figure 3B), whereas the adhesion proteins (E-cadherin, GEX2 and JUNO) did not display fusogenic activity; significantly, IZUMO1 induced fusion albeit to somewhat lower levels than EFF-1 (Figure 3B; Movies S2 and S3). Because IZUMO1 was expressed fused to Venus in the C terminus we could follow its expression; we often observed Venus fluorescence at time points when fusion started taking place (Figure 3A; Movies S2 and S3). Taken together, our results utilizing content mixing and live imaging support a model in which IZUMO1 acts independently of JUNO in a unilateral mechanism to fuse mammalian cells (Figure S1B).

**Figure 3.**
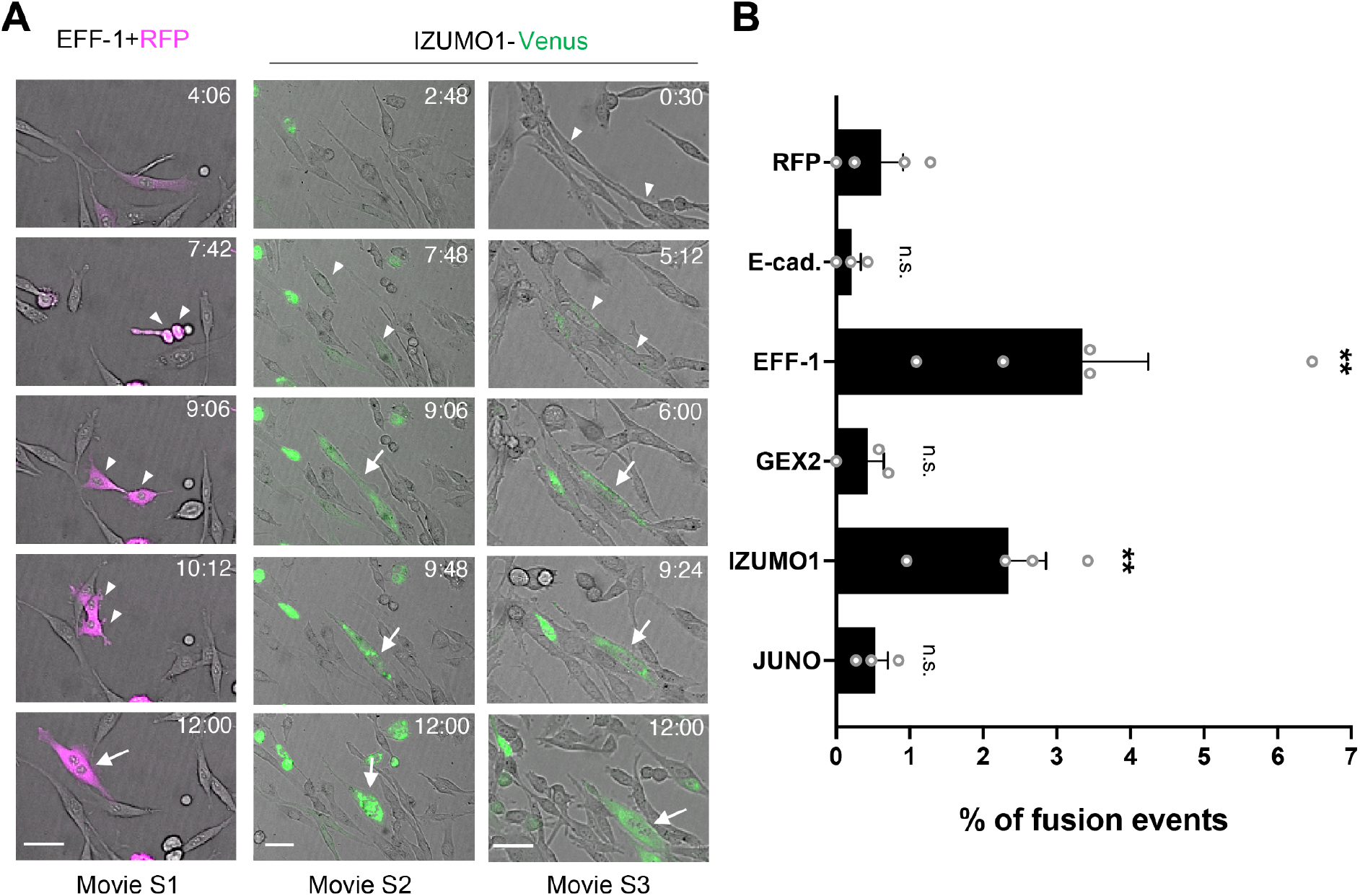
Quantification of fusion activity using live imaging. **(A)** Time-lapse images from a fusion assay. BHK cells were transfected with plasmids for expression of cytoplasmic RFP (RFPcyto, magenta) and EFF-1 (co-transfection) or IZUMO1-Venus (green). For IZUMO1-Venus, two independent fusion events are shown. Arrowheads and arrows indicate contacting and fused cells, respectively. Time (h:min) after the start of observation (see Movies S1-S3). Scale bars, 50 µm. **(B)** Quantification of live imaging experiments in which BHK cells express RFPcyto, E-cadherin, EFF-1, GEX2, IZUMO1-Venus or JUNO. The percentage of fusion was defined as the ratio between the number of fusion events (Fe) and the number of transfected cells (Tc), as follows: % of fusion events = (Fe/Tc) x 100. Bar chart showing individual experiments and means ± SEM of at least three independent experiments. Total Tc analyzed = 1001 (RFP), 1179 (E-cadherin), 930 (EFF-1), 415 (GEX2), 817 (IZUMO1) and 1265 (JUNO). Comparisons by one-way ANOVA followed by Dunett’s test against RFPcyto. ns = non-significant, * p < 0.05, ** p < 0.01.

### Somatic cells expressing IZUMO1 attach to mouse oocytes but fail to fuse

Following our observation that IZUMO1 can mediate fusion of heterologous somatic cells in culture, we then aimed to study a semi-heterologous system which retains the endogenous components of the oocyte, but not sperm, membranes. To this end, BHK cells expressing IZUMO1 along with the nuclear marker H2B-RFP were incubated with mouse oocytes expressing the transgenic protein CD9-GFP on their membrane and the occurrence of fusion was evaluated (Figure S1A). We found that IZUMO1 expression was sufficient to mediate the binding of BHK cells to oocytes but did not induce their fusion (Figure S4B), consistent with previous reports (Chalbi et al., 2014; Inoue et al., 2013, 2015). As a positive control, we used a powerful and promiscuous viral fusogen, the Vesicular Stomatitis Virus G protein (VSV-G) (Florkiewicz and Rose, 1984). We co-expressed IZUMO1 together with VSV-G to mediate BHK attachment to the oocytes, and lowered the pH to trigger VSV-G activity (Florkiewicz and Rose, 1984). Only in this condition we observed chromosomes from the BHK in the cytoplasm of the oocytes (Figure S4B). Thus, while viral VSV-G can efficiently fuse BHK cells to oocytes, we could not demonstrate a similar role for IZUMO1 under the conditions employed, which might suggest a role for additional cofactors that may be required to trigger fusion or bypass the mechanisms that block fusion to the egg.

### Dissection of the JUNO-binding and membrane fusion activities of IZUMO1

Some viral fusogens accomplish apposing membrane docking and elicit their merger using two different structural domains (White et al., 2008). We hypothesized that IZUMO1 will similarly have different domains mediating its dual activities. To study the structural features involved in IZUMO1 fusogenic activity, we designed and generated a series of mutants based on the crystal structure of IZUMO1 and its interactions with the docking receptor JUNO (Figures 4A and 4B). In order to assess the functionality of these mutants for the dual role of IZUMO1, we confirmed the expression of each of the mutant proteins by western blot (Figure 4C) and their localization by immunostaining (Figure 4D) and compared their performance in binding and content-mixing experiments. IZUMO1 contains the so-called Izumo domain composed of a four-helix bundle in the N-terminal part and a β-hairpin or hinge that links the Izumo domain to an Ig-like domain (Figure 4A; (Nishimura et al., 2016)). Single-pass transmembrane proteins with Ig-like domains have been involved in cell-to-cell fusion, like the members of the Fusexin superfamily (Martens and McMahon, 2008; Vance and Lee, 2020). We found that the deletion of the transmembrane domain produced an inactive form of the protein (IZUMO^Ecto^; Figure 5A) that was not detected on the surface of the cells, suggesting that anchoring to the plasma membrane is required for fusion. When the Ig-like domain of IZUMO1 was absent, the protein IZUMO1^ΔIg^ was expressed intracellularly (Figure 4C) but not detected on the cellular surface (Figure 4D), suggesting that this disruption interferes with the correct localization of the protein. While not detected on the cell surface using immunostaining, IZUMO1^ΔIg^ was able to induce significant levels of content-mixing compared to the empty vector, although lower than wild-type IZUMO1 (Figure 5A). These results suggest that some IZUMO1^ΔIg^ was able to reach the cell surface at low levels or transiently and was still able to induce fusion, detected as content mixing. Thus, IZUMO^Ecto^ cannot mediate cell-cell fusion suggesting that the transmembrane domain is necessary for fusion. In contrast, the Ig-like domain is required for correct trafficking and localization to the plasma membrane but does not appear to be essential for fusion, since IZUMO1^ΔIg^ is still able to mediate cell-cell fusion.

**Figure 4.**
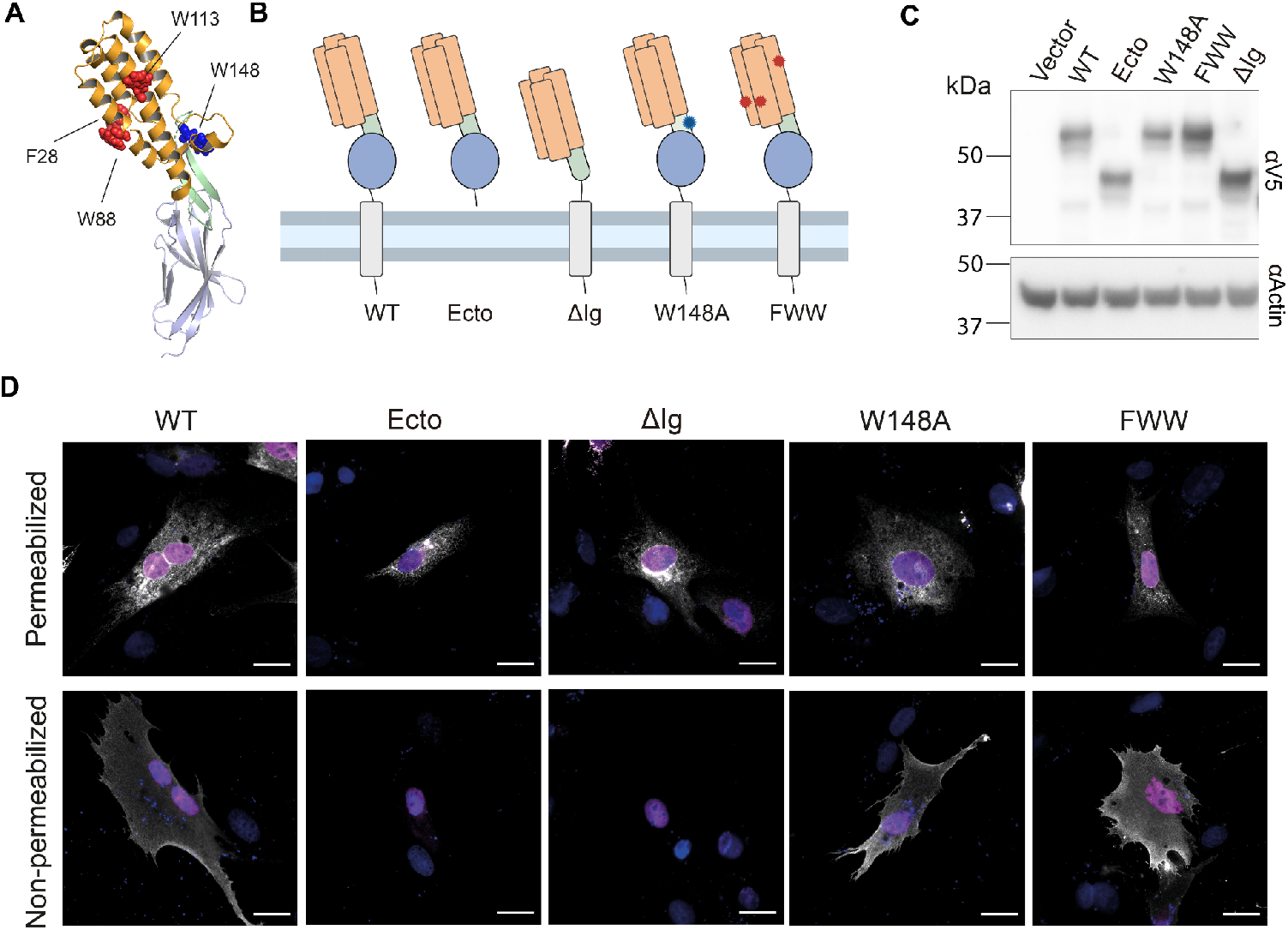
Mutagenesis of IZUMO1 and expression of mutant proteins. **(A)** Structure of IZUMO1 (PDB: 5B5K, (Nishimura et al., 2016)) showing the four-helix bundle (in orange) containing four solvent-exposed aromatic residues (in red, F28, W88 and W113); the hinge (in light green) with the JUNO-interacting W148 (in blue); and the Ig-like domain (in teal). **(B)** Schematic representation of wild-type IZUMO1 (WT) and the mutants maintaining the color coding for the different domains as in (A). The mutants with the deletion of the transmembrane domain and cytoplasmic tail (Ecto), the deletion of the Ig-like domain (ΔIg), the point mutation W148A and the triple mutant (F28A+W88A+W113A, FWW) are represented. **(C)** Representative Western blot of total protein extract from BHK cells transfected with empty vector, a plasmid encoding for WT IZUMO1, or the mutants shown in (B). The different variants were detected with an anti-V5 antibody and actin was used as a loading control. **(D)** Representative images of cells transfected with the pCI::H2B-RFP vectors encoding for WT IZUMO1 or the different mutants and subjected to an immunostaining (in white) after permeabilization with Triton X-100 using an anti-V5 antibody or without permeabilization using an antibody against the Izumo domain (Mab120). The signals for H2B-RFP and DAPI are shown in magenta and blue, respectively. The proportion of non-permeabilized cells showing surface expression was: IZUMO1^WT^ (41.9%, n=1013), IZUMO1^ΔIg^ (0%, n=1000), IZUMO1^Ecto^ (0%, n=1000), IZUMO1^W148A^ (60.3%, n=1005), IZUMO1^FWW^ (61.7%, n=995). Scale bars, 20 µm.

**Figure 5.**
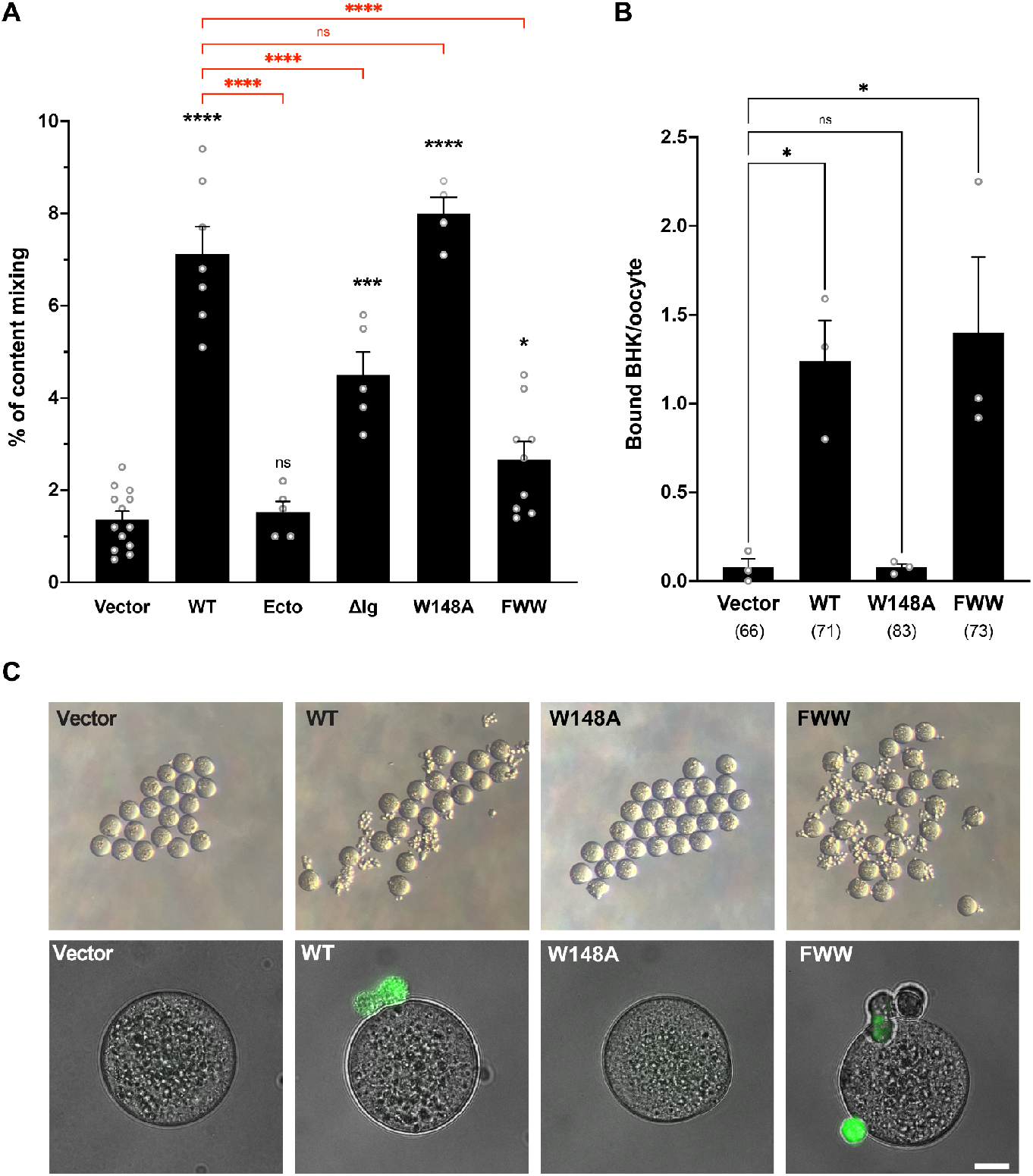
Functional characterization of IZUMO1 mutants. **(A)** Quantification of content-mixing in populations of cells expressing vectors, wild-type IZUMO1 (WT) and its mutants (Ecto, ΔIg, W148A and FWW, see Figure 4). Bar chart showing means ± SEM. n = 1000 nuclei per experiment. Comparisons by one-way ANOVA followed by Bonferroni’s test against the vector (black) and against WT (red). ns = non-significant, * p<0.05, *** p < 0.001, **** p< 0.0001. **(B-C)** Analysis of the binding ability of IZUMO1 (WT and mutants) to oocytes. **(B)** Quantification of the binding of BHK cells to oocytes. Cells were transfected with pCI::GFPnes empty vector or encoding for IZUMO1, IZUMO1^W148A^ or IZUMO1^FWW^ and incubated with wild-type oocytes. The number of BHK cells bound per oocyte was determined; n = total number of oocytes analyzed in parenthesis. Bar chart showing means ± SEM. Comparisons by one-way ANOVA followed by Dunnett’s test against the vector. ns = non-significant, * p<0.05. **(C)** Representative images of oocytes from one experiment in (**B**) taken under a dissecting microscope (upper row) and a wide-field illumination microscope showing the merged DIC and GFP channels (lower row). Scale bar, 20 µm.

To directly study whether the fusogenic activity of IZUMO1 is correlated with the ability to bind JUNO, we analyzed a mutant bearing a mutation in a conserved tryptophan residue in the hinge region that is essential for IZUMO1-JUNO interaction (W148A, Figure 4A, (Aydin et al., 2016; Ohto et al., 2016)). We found that IZUMO1^W148A^ had no effect on the levels of content mixing compared to wild-type IZUMO1 (Figure 5A) but disrupted the binding of BHK cells to oocytes (Figures 5B and 5C). Unlike IZUMO^Ecto^ or IZUMO1^ΔIg^, the W148A mutation did not affect its localization on the surface (Figure 4D). Thus, the W148A residue on the β-hairpin of IZUMO1 is required for binding the oocyte via JUNO but does not play an active role in cell-to-cell fusion.

The four-helix bundle of the Izumo domain contains three exposed aromatic residues (Figure 4A; (Nishimura et al., 2016)) which may be important for the interaction of IZUMO1 to the membrane of the oocyte or to other proteins given that JUNO is displaced from the fusion site shortly after the apposition of the fusing membranes (Inoue et al., 2015). To test whether these residues are required for efficient fusion, we generated a triple mutant F28A, W88A and W113A (FWW, Figure 4B). IZUMO1^FWW^ was detected on the surface of BHK cells (Figure 4D) and was still able to mediate BHK binding to the oocyte (Figures 5B and 5C), however, this mutant induced significantly lower levels of content mixing than wild-type IZUMO1 (Figure 5A). To summarize, our results support the existence of two functional domains in IZUMO1, with binding to JUNO mediated through W148 and cell-to-cell fusion dependent on three exposed aromatic residues on the four-helix bundle of the Izumo domain. These results allow us to uncouple the dual role of IZUMO1 as a JUNO-interacting binding protein and a unilateral fusogen, which utilize separate structural motifs.

## Discussion

### Aromatic residues on the four-helix bundle are required for fusion independently of the JUNO-IZUMO1 interactions

Our results show that IZUMO1 is sufficient to induce heterologous cell-to-cell fusion. While IZUMO1 shares structural similarities with proteins involved in cell invasion of malaria parasites (Nishimura et al., 2016), there is no obvious homology between IZUMO1 and any known fusogen. Additionally, IZUMO1 lacks any clear hydrophobic stretch that could work as a fusion peptide or loop which serves to anchor to the opposing membrane (Aydin et al., 2016), a characteristic feature which is shared by many, but not all, cellular and viral fusogens (Brukman et al., 2019). However, the four-helix bundle of the Izumo domain contains solvent-exposed aromatic residues (Nishimura et al., 2016); which are not involved in JUNO-IZUMO1 interaction (Aydin et al., 2016; Ohto et al., 2016) but might be relevant for interacting with the egg membrane. This is supported by our findings that the simultaneous mutation of these residues did not affect the transinteraction of IZUMO1-JUNO mediating binding of BHK cells to oocytes, but did reduce the fusogenic activity of IZUMO1 (Figure 5). While F28 is well conserved in rodents, W88 and W113 are present in most IZUMO1 mammalian orthologs (Aydin et al., 2016; Ohto et al., 2016). Interestingly, monoclonal antibodies against the N-terminal region of IZUMO1 were capable of inhibiting gamete fusion *in vitro* without altering sperm binding to the oocyte (Inoue et al., 2013), supporting not only an additional role of IZUMO1 in fusion besides binding, but also a relevance of the Izumo domain in these later stages of gamete fusion. Alternatively, the aromatic residues on the surface of the Izumo domain may be required for the interaction with an unknown secondary receptor (Inoue et al., 2015), for oligomerization (Ellerman et al., 2009) or for interaction with the other domains within IZUMO1 protein after a conformational change (Inoue et al., 2015).

### Spatial and temporal dissection of egg-sperm binding and fusion

Previous work has established that after sperm binding, JUNO is excluded from the interface between egg and sperm, while IZUMO1 is conversely enriched in this region (Inoue et al., 2015). These observations are consistent with a dual role of IZUMO1 as a fusogen. The concentration of IZUMO1 at the fusion zone is accompanied by a conformational change that depends on disulfide isomerase activity, and dimerization (Inoue et al., 2015). Furthermore, IZUMO1 has a more ancestral origin, being found in many vertebrate phyla, while JUNO is found in mammals only (Grayson, 2015). This evolutionary distribution, as well as IZUMO1’s unilateral fusogenic activity, suggest a function for IZUMO1 in fertilization that is independent of its well-established interaction with JUNO. This is further supported by our finding that the IZUMO1^W148A^ mutant which fails to bind JUNO is still able to induce cell-to-cell fusion, and that a triple mutant of the exposed aromatic residues in the four-helix bundle is able to mediate robust BHK binding to oocytes while cell–to-cell fusion was significantly reduced. Additionally, our results suggest a role for the IZUMO1 Ig-like domain for correct localization of the protein to the membrane (Figure 4D), however, it is not essential for its fusogenic activity (Figure 5A). This change in the localization could be explained by the lack of the single glycosylation site at the Ig-like domain that was reported to be relevant for protecting the protein from degradation but not for its activity in fertilization (Inoue et al., 2008).

### Are other egg and sperm membrane proteins involved in membrane merger?

While our results point to IZUMO1 as a *bona fide* fusogen that can fuse hamster and human cells in culture, this does not exclude the possibility that other fusogenic proteins work cooperatively with IZUMO1 to ensure the success and specificity of fertilization *in vivo* (Chalbi et al., 2014; Deneke and Pauli, 2021; Ellerman et al., 2009; Gaikwad et al., 2019; Inoue et al., 2013, 2015). This kind of cooperation can also be seen in plant fertilization, where the mutation of DMP8/9 proteins drastically reduces the activity of GCS1/HAP2 in vivo (Cyprys et al., 2019; Zhang et al., 2019). Proteins related to IZUMO1, for instance, IZUMO2-4, SPACA6 or TMEM95 may work as fusogens or potentiate IZUMO1’s fusogenic activity (Deneke and Pauli, 2021; Ellerman et al., 2009). Notably, IZUMO1 can homo- and hetero-oligomerize (Ellerman et al., 2009; Gaikwad et al., 2019). The absence of these proteins may explain the inability of somatic cells expressing IZUMO1 to fuse to oocytes (Figure S4B, also see (Chalbi et al., 2014; Inoue et al., 2013, 2015)). We cannot exclude other factors, such as the movement of the flagella (Ravaux et al., 2016) or the exposure of phosphatidylserine on the sperm (Rival et al., 2019), that may also be required for a successful fusion with the oocyte.

Additionally, the fact that other eukaryotic fusogens (*e*.*g*. EFF-1, AFF-1 and GCS1/HAP2) were shown to be less efficient *in vitro* than their viral counterparts (Avinoam et al., 2011; Valansi et al., 2017) might also contribute to explain why previous studies failed to detect fusion of somatic cells expressing IZUMO1 and JUNO (Bianchi et al., 2014).

### IZUMO1 activity is regulated during fertilization

Despite IZUMO1 being a unilateral fusogen, sperm cells do not fuse with other sperm or with other cell types in the male or female reproductive tracts. Furthermore, gamete fusion is species-specific (Yanagimachi, 1988). These non-physiological fusion events do not lead to a successful formation of a zygote and therefore reduce the individual fitness. The changes in the localization of IZUMO1 during sperm transit may partially explain this. In fresh sperm, IZUMO1 is not exposed on the outer membrane, being localized to the interior of the acrosome (Inoue et al., 2005), a giant specialized vesicle in the head, and therefore it cannot mediate cell-to-cell fusion. Only after capacitation and the acrosomal exocytosis in the female tract, IZUMO1 is exposed to the cell surface and migrates to the fusogenic region of the sperm plasma membrane (Inoue et al., 2005; Satouh et al., 2012). In addition, the requirement of a tight binding step to JUNO prior to fusion determines that the sperm cell will fuse to oocytes expressing a species-matching IZUMO1 receptor (Bianchi and Wright, 2015; Bianchi et al., 2014). This is a common mechanism used by many viruses to regulate their cellular tropism even when carrying powerful unilateral fusogens, such as the requirement of CD4 and other co-receptors for HIV infection mediated by the Env glycoprotein (Melikyan, 2011; White et al., 2008). Another analogy can be made with intracellular fusogens such as the SNARE complex that requires the regulatory activities of Munc18, Munc13 and synaptotagmin for efficient fusion (Stepien and Rizo, 2021). After sperm-to-oocyte fusion occurs, JUNO is shed from the plasma membrane preventing further sperm to bind, and therefore, contributing to a block to polyspermy (Bianchi et al., 2014).

### Concluding remarks

Up to this report, GCS1/HAP2 proteins were the only known fusogens involved in fertilization (reviewed in (Brukman et al., 2022)); these proteins are structurally and evolutionarily related to Class II fusogens from enveloped viruses and FF fusogens from nematodes (Fédry et al., 2017; Pinello et al., 2017; Valansi et al., 2017). Together, these fusogens form a superfamily called Fusexins. The involvement of GCS1/HAP2 in gamete fusion seems to be a characteristic of eukaryotes (Pinello et al., 2017; Valansi et al., 2017); however, many sexually reproducing organisms including fungi and vertebrates lack a fusexin homologue. Considering their distributions in the phylogenetic tree (Vance and Lee, 2020), it is possible that IZUMO1-type proteins are chordate’s innovations that replaced GCS1/HAP2 during evolution as a strategy for gamete fusion. Combined with previous information (Deneke and Pauli, 2021; Ellerman et al., 2009; Inoue et al., 2013, 2015)(Stepien and Rizo, 2021), we hypothesize that IZUMO1 from the sperm first transiently binds JUNO on the oocyte for docking and subsequently undergoes a conformational change and oligomerization that induce sperm-egg fusion (Figure S1B). This role in membrane merging makes IZUMO1 even more suited to its name, coined after the Japanese shrine dedicated to marriage (Inoue et al., 2005).

## Acknowledgments

We thank Gavin Wright for the plasmids containing *mIzumo1* and *mJuno* sequences and Masahito Ikawa for the CD9-GFP transgenic mouse line. We thank Pablo Aguilar, Dan Cassel, Masahito Ikawa, Yael Iosilevskii and Luca Jovine for critically reading the manuscript. This work was funded by the Israel Science Foundation (ISF grants 257/17, 2462/18, 2327/19 and 178/20 to B.P.), by Grant-in-Aid for Scientific Research on Innovative Areas (16H06465, 16H06464, and 16K21727 to T.H.), by JST, ERATO (JPMJER1004 to T.H.) and CREST (JPMJCR20E5 to T.H.). This project has received funding from the European Union’s Horizon 2020 research and innovation programme under the Marie Skłodowska-Curie (grant agreement No 844807 to N.G.B).

## Author contributions

B.P. and T.H. conceived the study. N.G.B., K.P.N., C.V., X.L. and K.F. designed and conducted experiments, and processed and analyzed the data supervised by B.P. and T.H.; N.G.B. and B.P. wrote the initial draft of the manuscript. All authors participated in discussions of results and manuscript editing.

## STAR★ Methods

### KEY RESOURCES TABLE

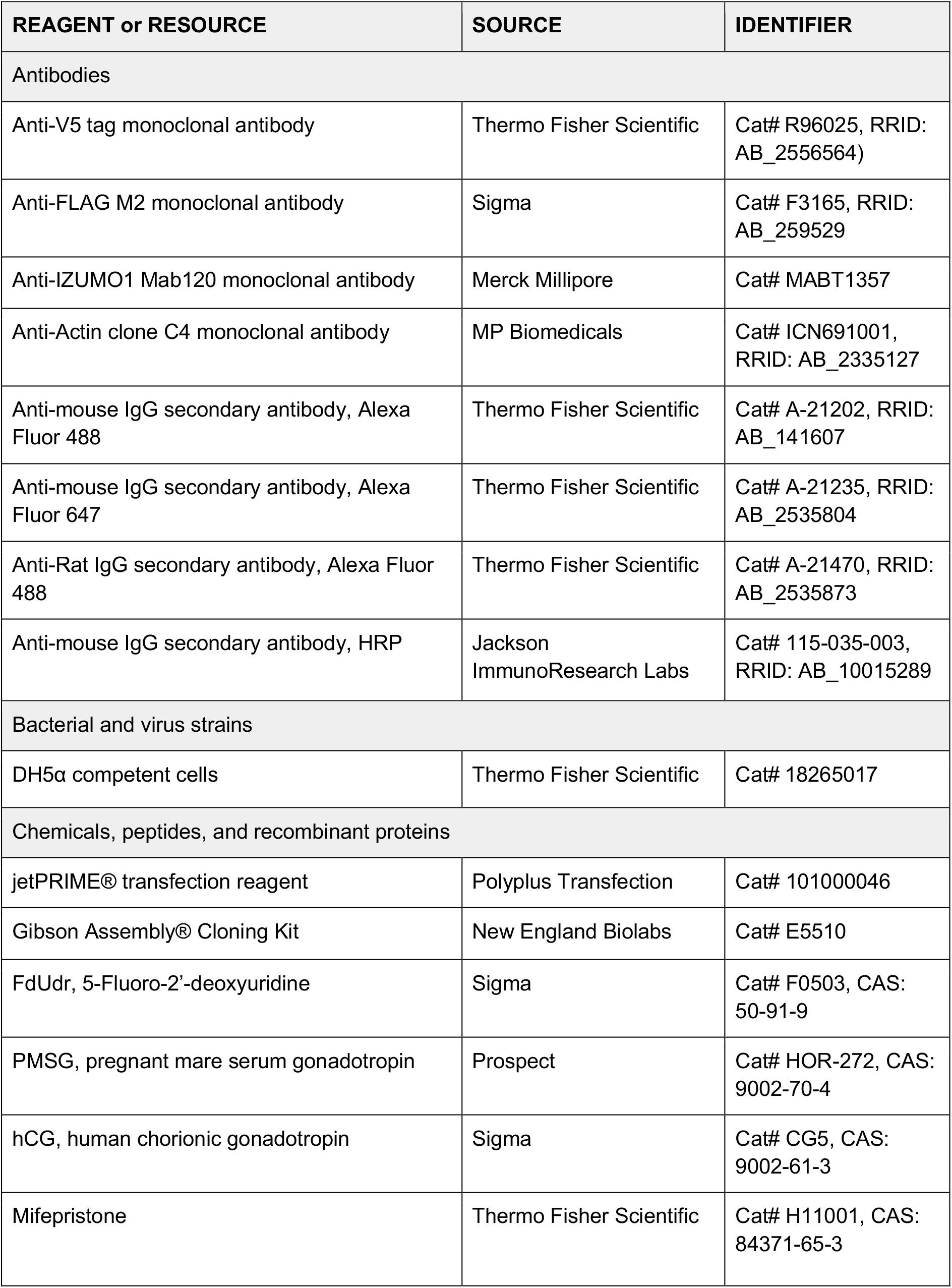
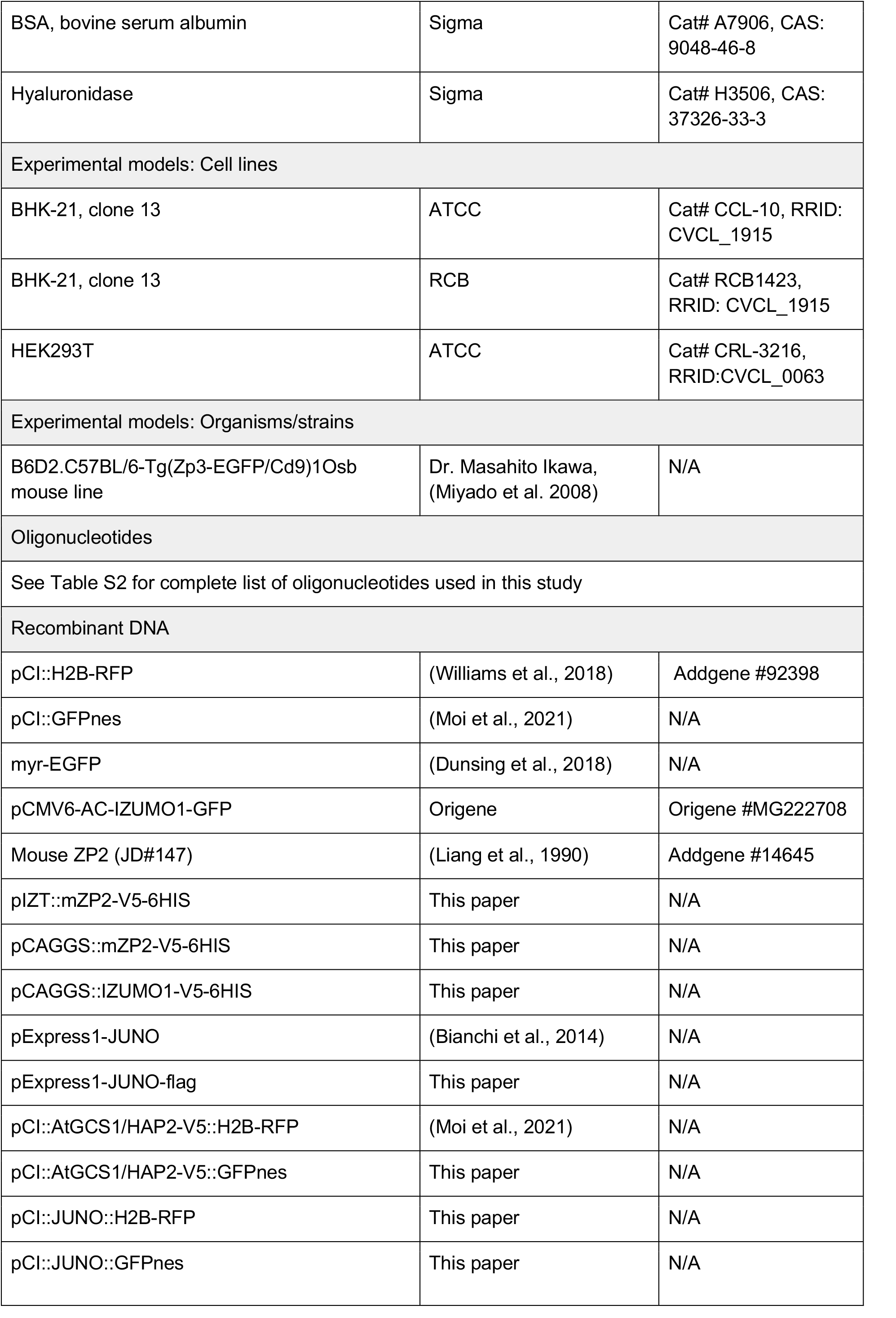
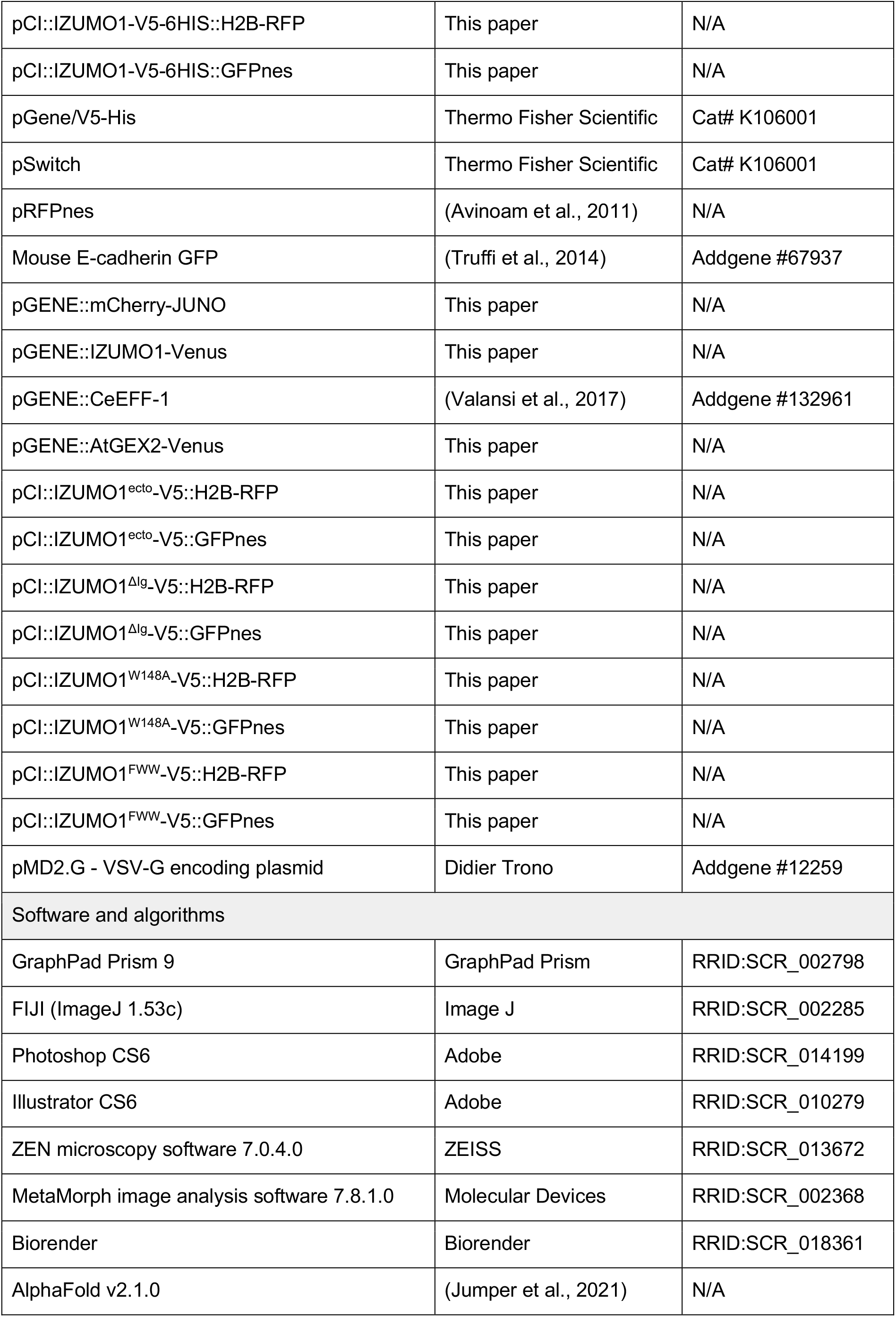

| REAGENT or RESOURCE | SOURCE | IDENTIFIER |
| --- | --- | --- |
| Antibodies |  |  |
| Anti-V5 tag monoclonal antibody | Thermo Fisher Scientific | Cat# R96025, RRID: AB_2556564) |
| Anti-FLAG M2 monoclonal antibody | Sigma | Cat# F3165, RRID: AB_259529 |
| Anti-IZUMO1 Mab120 monoclonal antibody | Merck Millipore | Cat# MABT1357 |
| Anti-Actin clone C4 monoclonal antibody | MP Biomedicals | Cat# ICN691001, RRID: AB_2335127 |
| Anti-mouse IgG secondary antibody, Alexa Fluor 488 | Thermo Fisher Scientific | Cat# A-21202, RRID: AB_141607 |
| Anti-mouse IgG secondary antibody, Alexa Fluor 647 | Thermo Fisher Scientific | Cat# A-21235, RRID: AB_2535804 |
| Anti-Rat IgG secondary antibody, Alexa Fluor 488 | Thermo Fisher Scientific | Cat# A-21470, RRID: AB_2535873 |
| Anti-mouse IgG secondary antibody, HRP | Jackson ImmunoResearch Labs | Cat# 115-035-003, RRID: AB_10015289 |
| Bacterial and virus strains |  |  |
| DH5α competent cells | Thermo Fisher Scientific | Cat# 18265017 |
| Chemicals, peptides, and recombinant proteins |  |  |
| jetPRIME® transfection reagent | Polyplus Transfection | Cat# 101000046 |
| Gibson Assembly® Cloning Kit | New England Biolabs | Cat# E5510 |
| FdUdr, 5-Fluoro-2'-deoxyuridine | Sigma | Cat# F0503, CAS: 50-91-9 |
| PMSG, pregnant mare serum gonadotropin | Prospect | Cat# HOR-272, CAS: 9002-70-4 |
| hCG, human chorionic gonadotropin | Sigma | Cat# CG5, CAS: 9002-61-3 |
| Mifepristone | Thermo Fisher Scientific | Cat# H11001, CAS: 84371-65-3 |
| BSA, bovine serum albumin | Sigma | Cat# A7906, CAS: 9048-46-8 |
| Hyaluronidase | Sigma | Cat# H3506, CAS: 37326-33-3 |
| Experimental models: Cell lines |  |  |
| BHK-21, clone 13 | ATCC | Cat# CCL-10, RRID: CVCL_1915 |
| BHK-21, clone 13 | RCB | Cat# RCB1423, RRID: CVCL_1915 |
| HEK293T | ATCC | Cat# CRL-3216, RRID: CVCL_0063 |
| Experimental models: Organisms/strains |  |  |
| B6D2.C57BL/6-Tg(Zp3-EGFP/Cd9)1Osb mouse line | Dr. Masahito Ikawa, (Miyado et al. 2008) | N/A |
| Oligonucleotides |  |  |
| See Table S2 for complete list of oligonucleotides used in this study |  |  |
| Recombinant DNA |  |  |
| pCI::H2B-RFP | (Williams et al., 2018) | Addgene #92398 |
| pCI::GFPnes | (Moi et al., 2021) | N/A |
| myr-EGFP | (Dunsing et al., 2018) | N/A |
| pCMV6-AC-IZUMO1-GFP | Origene | Origene #MG222708 |
| Mouse ZP2 (JD#147) | (Liang et al., 1990) | Addgene #14645 |
| pIZT::mZP2-V5-6HIS | This paper | N/A |
| pCAGGS::mZP2-V5-6HIS | This paper | N/A |
| pCAGGS::IZUMO1-V5-6HIS | This paper | N/A |
| pExpress1-JUNO | (Bianchi et al., 2014) | N/A |
| pExpress1-JUNO-flag | This paper | N/A |
| pCI::AtGCS1/HAP2-V5::H2B-RFP | (Moi et al., 2021) | N/A |
| pCI::AtGCS1/HAP2-V5::GFPnes | This paper | N/A |
| pCI::JUNO::H2B-RFP | This paper | N/A |
| pCI::JUNO::GFPnes | This paper | N/A |
| pCI::IZUMO1-V5-6HIS::H2B-RFP | This paper | N/A |
| pCI::IZUMO1-V5-6HIS::GFPnes | This paper | N/A |
| pGene/V5-His | Thermo Fisher Scientific | Cat# K106001 |
| pSwitch | Thermo Fisher Scientific | Cat# K106001 |
| pRFPnes | (Avinoam et al., 2011) | N/A |
| Mouse E-cadherin GFP | (Truffi et al., 2014) | Addgene #67937 |
| pGENE::mCherry-JUNO | This paper | N/A |
| pGENE::IZUMO1-Venus | This paper | N/A |
| pGENE::CeEFF-1 | (Valansi et al., 2017) | Addgene #132961 |
| pGENE::AtGEX2-Venus | This paper | N/A |
| pCI::IZUMO1<sup>ecto</sup>-V5::H2B-RFP | This paper | N/A |
| pCI::IZUMO1<sup>ecto</sup>-V5::GFPnes | This paper | N/A |
| pCI::IZUMO1<sup>Δlg</sup>-V5::H2B-RFP | This paper | N/A |
| pCI::IZUMO1<sup>Δlg</sup>-V5::GFPnes | This paper | N/A |
| pCI::IZUMO1<sup>W148A</sup>-V5::H2B-RFP | This paper | N/A |
| pCI::IZUMO1<sup>W148A</sup>-V5::GFPnes | This paper | N/A |
| pCI::IZUMO1<sup>FWW</sup>-V5::H2B-RFP | This paper | N/A |
| pCI::IZUMO1<sup>FWW</sup>-V5::GFPnes | This paper | N/A |
| pMD2.G - VSV-G encoding plasmid | Didier Trono | Addgene #12259 |
| Software and algorithms |  |  |
| GraphPad Prism 9 | GraphPad Prism | RRID:SCR_002798 |
| FIJI (ImageJ 1.53c) | Image J | RRID:SCR_002285 |
| Photoshop CS6 | Adobe | RRID:SCR_014199 |
| Illustrator CS6 | Adobe | RRID:SCR_010279 |
| ZEN microscopy software 7.0.4.0 | ZEISS | RRID:SCR_013672 |
| MetaMorph image analysis software 7.8.1.0 | Molecular Devices | RRID:SCR_002368 |
| Biorender | Biorender | RRID:SCR_018361 |
| AlphaFold v2.1.0 | (Jumper et al., 2021) | N/A |

### RESOURCE AVAILABILITY

#### Lead contact

Further information and requests for resources and reagents should be directed to and will be fulfilled by the lead contact, Benjamin Podbilewicz.

#### Data and code availability

Any additional information required to reanalyze the data reported in this paper is available from the lead contact upon request.

### EXPERIMENTAL MODEL AND SUBJECT DETAILS

#### Cell lines and DNA transfection

In this study we used BHK (CCL-10; ATCC, Virginia, USA) for multinucleation, content mixing and BHK to oocyte interaction experiments and for evaluation of expression by Western blot and immunostaining; BHK (RCB1423; RIKEN Cell Bank, Tsukuba, Japan), for live imaging assays and HEK293T (CRL-3216; ATCC) cells for content mixing experiments. BHK and HEK293T cells were grown and maintained in Dulbecco’s modified Eagle’s medium containing 10% fetal bovine serum (FBS). Cells were cultured at 37°C in 5% CO_2_. Plasmids were transfected into cells using 2 µl jetPRIME (PolyPlus-transfection, Illkirch-Graffenstaden, France) per µg of DNA in 100 µl of reaction buffer for every ml of medium. For experiments with HEK293T cells, a coating with poly-L-lysine hydrobromide (Sigma, 20 µg/ml) was applied to the plates.

#### Mice

All animal studies were approved by the Committee on the Ethics of Animal Experiments of the Technion, Israel institute of Technology. B6D2.C57BL/6-Tg(Zp3-EGFP/Cd9)1Osb mice line (Miyado et al., 2008) was obtained from Dr. Masahito Ikawa (Osaka University, Japan) and animals were bred and housed in the Technion animal facility under specific pathogen-free conditions with *ad libitum* access to food and water. The primers used for genotyping are outlined in Table S2. Transgenic and wild type females between 2-6 months were used for the experiments.

### METHOD DETAILS

#### DNA constructs

For the multinucleation and content mixing assays: Mouse *Izumo1* and *Juno* coding sequences were amplified from pCMV6-IZUMO1-GFP (MG222708, Origene, Rockville, MD, USA) and pExpress1-JUNO (Clone B2) plasmids, respectively, kindly provided by Gavin Wright. The GFP tag of IZUMO1 construct was replaced with V5-HIS tags during cloning. *Izumo1-V5* and *Juno* sequences, and *Arabidopsis thaliana GCS1HAP2*-V5 were subcloned by restriction cloning into pCI::H2B-RFP and pCI::GFPnes vectors separately employing enzymes from Thermo Fisher Scientific (Invitrogen, Waltham, MA, USA). These bicistronic vectors translate for a nuclear RFP (H2B-RFP) or cytoplasmic GFP (GFPnes) after an IRES element (Internal Ribosome Entry Site). For mutagenesis of IZUMO1: i) IZUMO1^ecto^: the ectodomain of IZUMO1 (M1-P312) was amplified; ii) IZUMO1^ΔIg^: the upstream and downstream of Ig domain (G166-L253) were amplified independently from the full length IZUMO1 and fused together by overlap PCR; iii) IZUMO1^W148A^: The W148 was mutated to Alanine by overlapping PCR; iv) IZUMO1^FWW^: F28, W88 and W113 located in the Izumo domain were mutated to Alanines by overlapping primers and fused together. The folding of the mutants was corroborated by AlphaFold (Jumper et al., 2021). All mutants were ligated into pCI::H2B-RFP and pCI::GFPnes vectors for mixing assay. JUNO was tagged in the C-terminus by inserting an annealed oligo containing the FLAG sequence after the signal peptide into the BlpI restriction site. Oligonucleotides were obtained from Sigma-Aldrich or IDT and all constructs were verified by DNA sequencing (Macrogen). For the live imaging experiments: *Arabidopsis thaliana GEX2, Caenorhabditis elegans eff-1* and mouse *Izumo* or *Juno* sequences were amplified from cDNAs. To visualize the proteins, the fragments corresponding to *GEX2* and *Izumo1* were fused to fluorescent protein sequences by PCR and then cloned into pGENE B after double digestion with restriction enzymes using the Gibson assembly (NEB, #E5510, MA, USA). pGENE::mCherry-JUNO was made by ligating mCherry and pGENE::JUNO fragments (Nakajima et al., 2021). For inducible expression using mifepristone in BHK cells, we used the GeneSwitch System (Invitrogen). The complete lists of plasmids and primers used in this study are shown in Key Resource Table and Table S2, respectively.

#### Immunostaining and analysis of localization

BHK cells were grown on 24-well glass bottom tissue-culture plates or in tissue-culture plates with coverslips. 24 h after plating, cells were transfected and incubated for additional 24 h before proceeding to the immunostaining. For IZUMO1 (WT) and its mutants (IZUMO1^ecto^, IZUMO1^ΔIg^, IZUMO1^W148A^ and IZUMO1^FWW^) encoded in the pCI::H2B-RFP vector cells were fixed with 4% PFA in PBS and, when indicated, permeabilized with 0.1% Triton X-100 in PBS. To detect the proteins immunofluorescence was performed with anti-V5 (1:500, R96025, Thermo Fisher Scientific) or anti-IZUMO1, clone Mab120 (1:500, MABT1357, Merck Millipore) followed by the secondary antibodies Alexa Fluor 488 goat anti-mouse (1:500, A21202, Thermo Fisher Scientific) and Alexa Fluor 488 chicken anti-rat (1:500, A21470, Thermo Fisher Scientific), respectively. For JUNO localization, pExpress1-JUNO-flag plasmid was transfected together with the empty pCI::H2B-RFP and the incubation with the first antibody anti-flag (1:1000, F3165, Sigma) was performed before fixation (for detecting surface protein) or after fixation and permeabilization (for detecting total protein). Then, JUNO was probed using the secondary Alexa Fluor 488 goat anti-mouse. In all cases, the nuclei were stained with 1 µg/ml DAPI and micrographs were obtained using wide-field illumination using an ELYRA system S.1 microscope (Plan-Apochromat 20X NA 0.8; Zeiss).

#### Evaluation of multinucleation

BHK cells were grown on 24-well glass bottom tissue-culture plates. 24 h after plating, cells were transfected with pCI::myrGFP::H2B-RFP (myristoylated EGFP), pCI::GCS1/HAP-V5::H2B-RFP, pCI::IZUMO1-V5:H2B-RFP or pCI::JUNO::H2B-RFP vectors encoding for myristoylated EGFP (myrGFP, gray), GCS1/HAP2-V5, IZUMO1-V5 or JUNO. For JUNO a plasmid for myrGFP (gray) was co-transfected. 24 h post-transfection, 20 µM 5-fluoro-2’-deoxyuridine (FdUrd) was added to the plates to arrest the cell cycle and 24 h later, the cells were fixed with 4% PFA in PBS and processed for immunofluorescence using an anti-V5 antibody, as explained before. Micrographs were obtained using wide-field illumination using an ELYRA system S.1 microscope (Plan-Apochromat 20X NA 0.8; Zeiss). Multinucleation percentage (Figure S1A) was determined as the ratio between the number of nuclei in multinucleated cells (NuM) and the total number of nuclei in multinucleated cells and expressing cells that were in contact but did not fuse (NuC) as follows: (Num/(Nuc + Num)) × 100.

#### Content mixing experiments

BHK or HEK293T cells at 70% confluence in 35 mm plates were transfected with 1 µg pCI::H2B-RFP or pCI::GFPnes (empty vectors as negative controls); pCI::GCS1/HAP2-V5::H2B-RFP or pCI::GCS1/HAP2-V5::GFPnes (positive control); pCI::IZUMO1-V5::H2B-RFP or pCI::IZUMO1-V5::GFPnes; pCI::JUNO::H2B-RFP or pCI::JUNO::GFPnes. 4 h after transfection, the cells were washed 4 times with DMEM with 10% serum, 4 times with PBS and detached using Trypsin (Biological Industries). The transfected cells were collected, resuspended in DMEM with 10% serum, and counted. Equal amounts of H2B-RFP and GFPnes cells (1-1.25×10^5^ each) were mixed and seeded on glass-bottom plates (12-well black, glass-bottom #1.5H; Cellvis) and incubated at 37°C and 5% CO_2_. For IZUMO1, pCI::IZUMO-V5::GFPnes cells were also mixed with pCI::H2B-RFP or pCI::JUNO::H2B-RFP transfected cells. 18 h after mixing, 20 µM FdUrd was added to the BHK cells. The mixed cells were co-incubated for a total of 48 h after which they were fixed with 4% PFA in PBS and stained with 1 µg/ml DAPI. Micrographs were obtained using wide-field illumination using an ELYRA system S.1 microscope (Plan-Apochromat 20X NA 0.8; Zeiss). The percentage of mixing was defined as the ratio between the nuclei in mixed cells (NuM) and the total number of nuclei in mixed cells and fluorescent cells in contact that did not fuse (NuC), as follows: % of mixing = (NuM/(NuM+NuC)) x 100 (Figure S1A). 1000 nuclei (NuM+NuC) were counted in each independent repetition (experimental point). For immunostaining after the content mixing, cells were treated as explained above, using as the secondary antibody the Alexa Fluor 647 goat anti-mouse (A21235, Thermo Fisher Scientific).

#### Live imaging experiments

To evaluate fusion by live imaging, we transfected BHK cells with pGENE and pSWITCH. 24 h after transfection, BHK cells were cultured at 5.0 × 10^4^ cells/ml. 4 h after transfection, the expression was induced by addition of 10−^4^ mM mifepristone. 3–4 h post-induction, images of the cells were acquired every 6 min for 12 h to record cell-to-cell fusion, using a spinning disk confocal system (CellVoyager CV1000; Yokogawa Electric, Tokyo, Japan) at a magnification of 10X (NA 0.40, 10×UPLSAPO; Olympus, Tokyo, Japan) dry objective. The number of transfected cells and the occurrence of fusion were evaluated. Image analyses were performed using CV1000 software (Yokogawa Electric) and the FIJI online tool was used to adjust the brightness and contrast. The percentage of fusion was defined as the ratio between the number of fusion events (Fe) and the number of transfected cells (Tc), as follows: % of fusion = Fe/Tc (Figure S1A).

#### Western blots

BHK cells at 70% confluence in 35 mm plates were transfected with 1 µg pCI::GFPnes (empty plasmid as negative controls); pCI::IZUMO1-V5::GFPnes; pCI::IZUMO1^Ecto^-V5::GFPnes, pCI::IZUMO1^ΔIg^-V5::GFPnes, pCI::IZUMO1^W148A^-V5::GFPnes or pCI::IZUMO1^FWW^-V5::GFPnes. 24 h post-transfection, cells were mixed with reducing sample buffer (#S3401, Sigma) and incubated 5 min at 95°C. Samples were loaded on a 10% SDS-PAGE gel and transferred to PVDF membrane. After blocking, membranes were incubated with primary antibody anti–V5 mouse monoclonal antibody (1:5,000) or anti-actin (1:2,000, ICN691001, MP Biomedicals) for 1 h at room temperature and HRP-conjugated goat anti-mouse secondary antibody (1:10,000, 115-035-003, Jackson ImmunoResearch Labs) 1 h at room temperature. Membranes were imaged by the ECL detection system using FUSION-PULSE.6 (VILBER).

#### Fusion of BHK cells to oocytes

BHK cells at 70% confluence in 35 mm plates were transfected with 1 µg pCI::IZUMO1::H2B-RFP alone or together with pMD2.G (encoding for VSV-G) 24 h before the collection of the oocytes. To induce ovulation, transgenic females were treated with an i.p. injection of pregnant mare serum gonadotropin (PMSG; 5 IU; #HOR-272, Prospec, Israel). followed by an i.p. injection of human chorionic gonadotropin (hCG; 5 IU, #CG5, Sigma) 48 h later. Cumulus-oocyte complexes were collected from the ampullae of induced females 12–15 h post-hCG administration in mHTF medium (Kito et al., 2004). The oocytes were denuded from the cumulus and the ZP by sequential treatment with 0.3 mg/ml hyaluronidase (H3506; Sigma) and acid Tyrode solution (pH 2.5) (Nicolson et al., 1975). BHK cells were harvested with 0.05% EDTA in PBS and washed once with mHTF. 15-20 ZP-free oocytes were incubated with 5×10^4^ BHK cells for 15 min with occasional mixing and washed. In the case of VSV-G-expressing cells, the oocytes were treated for 30 s with acid Tyrode solution (pH 2.5) and washed again with mHTF. In all cases, the oocytes-cells complexes were incubated for 5 h at 37°C and 5% CO_2_. Oocytes were then fixed, stained with DAPI and evaluated for the presence of H2B-RFP chromosomes within the cytoplasm (Figure S1A), using a CSU-X1 spinning disk confocal (Yokogawa) on a Nikon Eclipse Ti inverted microscope (Plan-Apochromat 60X NA 1.4, Nikon). Images were obtained using an iXon3 EMCCD camera (ANDOR) through MetaMorph (Molecular Devices, version 7.8.1.0).

#### Binding of BHK cells to oocytes

BHK cells at 70% confluence in 35 mm plates were transfected with 1 µg pCI::GFPnes, pCI::IZUMO1::GFPnes, pCI::IZUMO1^W148A^::GFPnes or pCI::IZUMO1^FWW^::GFPnes 24 h before the collection of the oocytes. Oocytes were obtained from wild type females as described above, incubated with the 5×10^4^ BHK cells with occasional mixing for 15 min, washed and fixed. The average number of BHK in direct contact with the oocyte was determined for each group. Micrographs were obtained using wide-field illumination as described above.

### QUANTIFICATION AND STATISTICAL ANALYSIS

#### Statistics and data analysis

Results are presented as means ± SEM. For each experiment we performed at least three independent biological repetitions. To evaluate the significance of differences between the averages we used one-way ANOVA as described in the legends (GraphPad Prism 9). Figures were prepared with Photoshop CS6 and Illustrator CS6 (Adobe), BioRender.com and FIJI (ImageJ 1.53c).

