## Supplementary Information for "IZUMO1 is a sperm fusogen"

This document contains:
Figures S1-S5
Tables S1 and S2
Movies Legends S1-S3

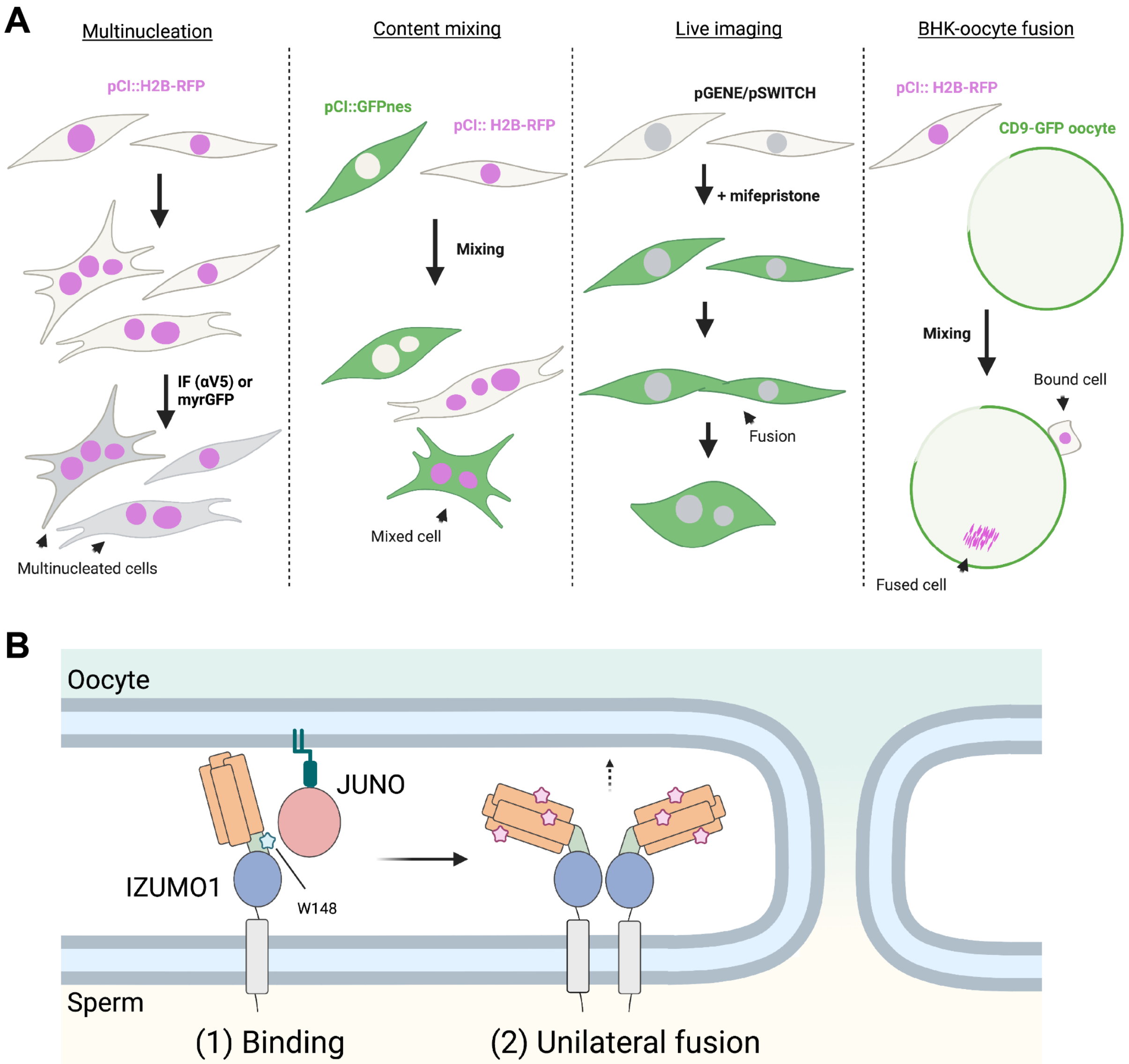


**Figure S1. Experimental designs and working model, related to Figures 1-5**
**(A)** Schemes of experimental designs.

*Multinucleation*: BHK cells are transfected with pCI::myrGFP::H2B-RFP (myristoylated GFP), pCI::GCS1/HAP-V5::H2B-RFP, pCI::IZUMO1-V5:H2B-RFP or pCI::JUNO::H2B-RFP. For JUNO, a plasmid for myrGFP was co-transfected, and the formation of cells with more than one nucleus is quantified. The outline of the cells is observed by performing immunostaining (IF) with antibodies against the V5 tag (**α**V5) or by detection of myristoylated GFP (myrGFP) fluorescence.

*Content mixing*: Cells (BHK or HEK293T) were transfected in two subpopulations with either a pCI::H2B-RFP or pCI::GFPnes plasmid for each candidate and mixed. The appearance of mixed cells containing both fluorescent markers was evaluated in relation to fluorescent cells which contact one another but are not content-mixed.

*Live imaging*: BHK cells were transfected with plasmids of the inducible system pGENE/pSWITCH encoding for each candidate fusogen. For those candidates without a fluorescent tag, a plasmid encoding for cytoplasmic RFP was co-transfected. The candidate proteins were induced by mifepristone and the cells subsequently tracked using live imaging. The occurrence of fusion was registered and quantified. The signal corresponding to the fluorescent tags or cytoplasmic RFP was also detected.

*BHK-oocyte fusion*: BHK cells were transfected with pCI::IZUMO1-V5:H2B-RFP alone or together with a plasmid encoding for VSV-G and mixed with ZP-free oocytes expressing CD9-GFP on the fusogenic region of the membrane. The presence of RFP-positive chromosomes within the cytoplasm of the oocytes was evaluated.

**(B)** Working model for IZUMO1 activities. 1. Transient interaction between JUNO and IZUMO1 (“Binding”) mediated by IZUMO1 W148 (blue star). 2. Conformational change of IZUMO1, oligomerization (*13*) and unilateral induction of gamete fusion (“Unilateral fusion”), enhanced by the action of F28, W88 and W113, the surface exposed aromatic residues in the four-helix bundle of IZUMO1 (pink stars). IZUMO1 domain colors correspond to Figures 4A and 4B.

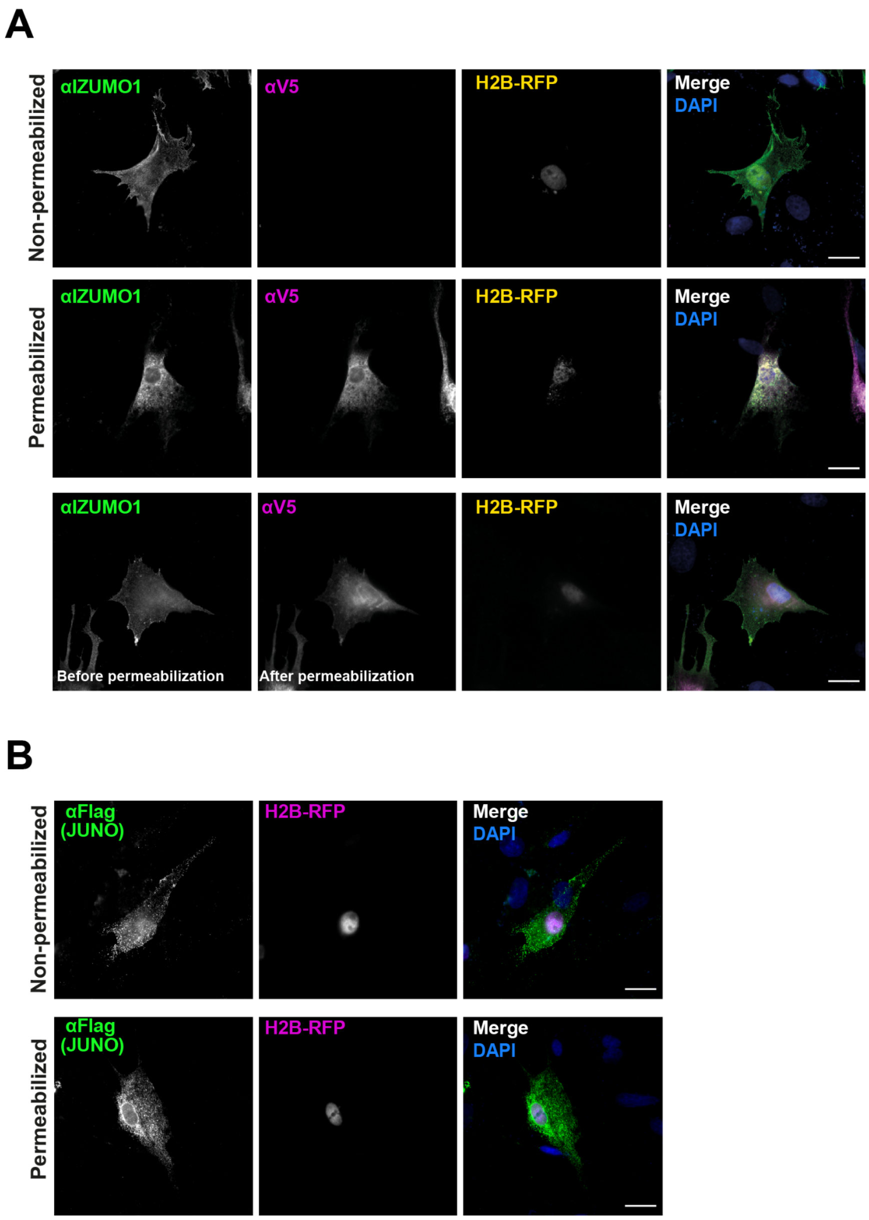


**Figure S2. Surface localization of IZUMO1 and JUNO, related to Figure 1**

**(A)** IZUMO1 presence on the surface of BHK cells transfected with pCI::IZUMO1-V5::H2B-RFP was determined by immunostaining using an anti-IZUMO1 (Mab 120) that recognizes the extracellular N-terminal region of the protein (green) or anti-V5 against the cytoplasmic C-terminal tag (magenta). Non-permeabilized and permeabilized cells were tested. Also, cells were exposed to anti-IZUMO1, then permeabilized and finally incubated with anti-V5 (lower row). In all cases, DAPI staining is shown in blue in the merge. Scale bars, 20 µm.

**(B)** JUNO localization to the plasma membrane of BHK transfected with pExpress1-JUNO-flag and pCI::H2B-RFP was evaluated by incubating the cells with anti-flag antibody before fixation (non-permeabilized) or after fixation and permeabilization. The immunostaining signal (green) and the nuclei of transfected cells (magenta) are shown in separate channels and in the merge which includes the DAPI staining (blue). Scale bars, 20 µm.

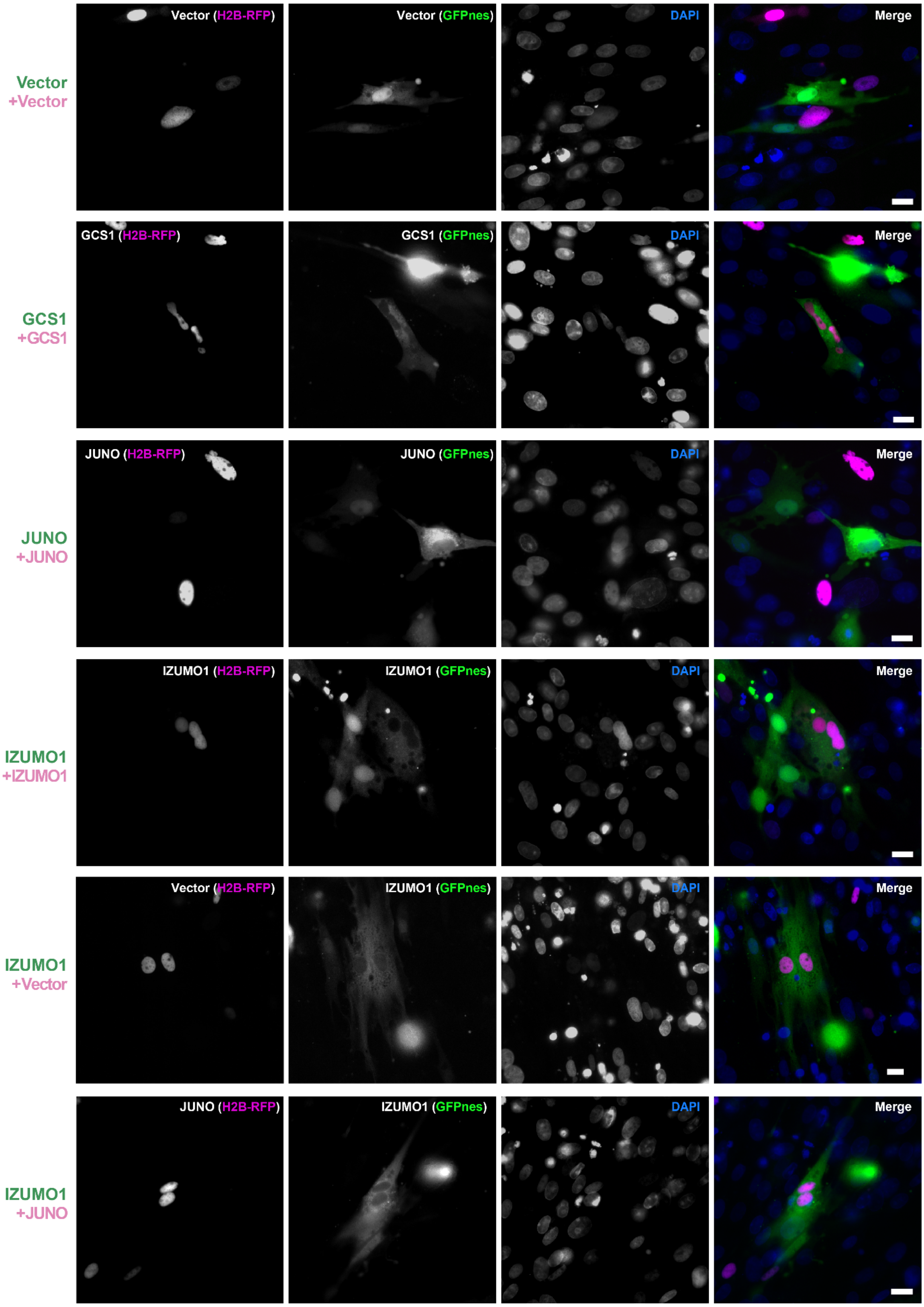


**Figure S3. Evaluation of content mixing of BHK cells, related to Figure 2**
Images from **Figure 2A** in each separate channel (red, green and DAPI) and merge. Scale bars, 20 µm.

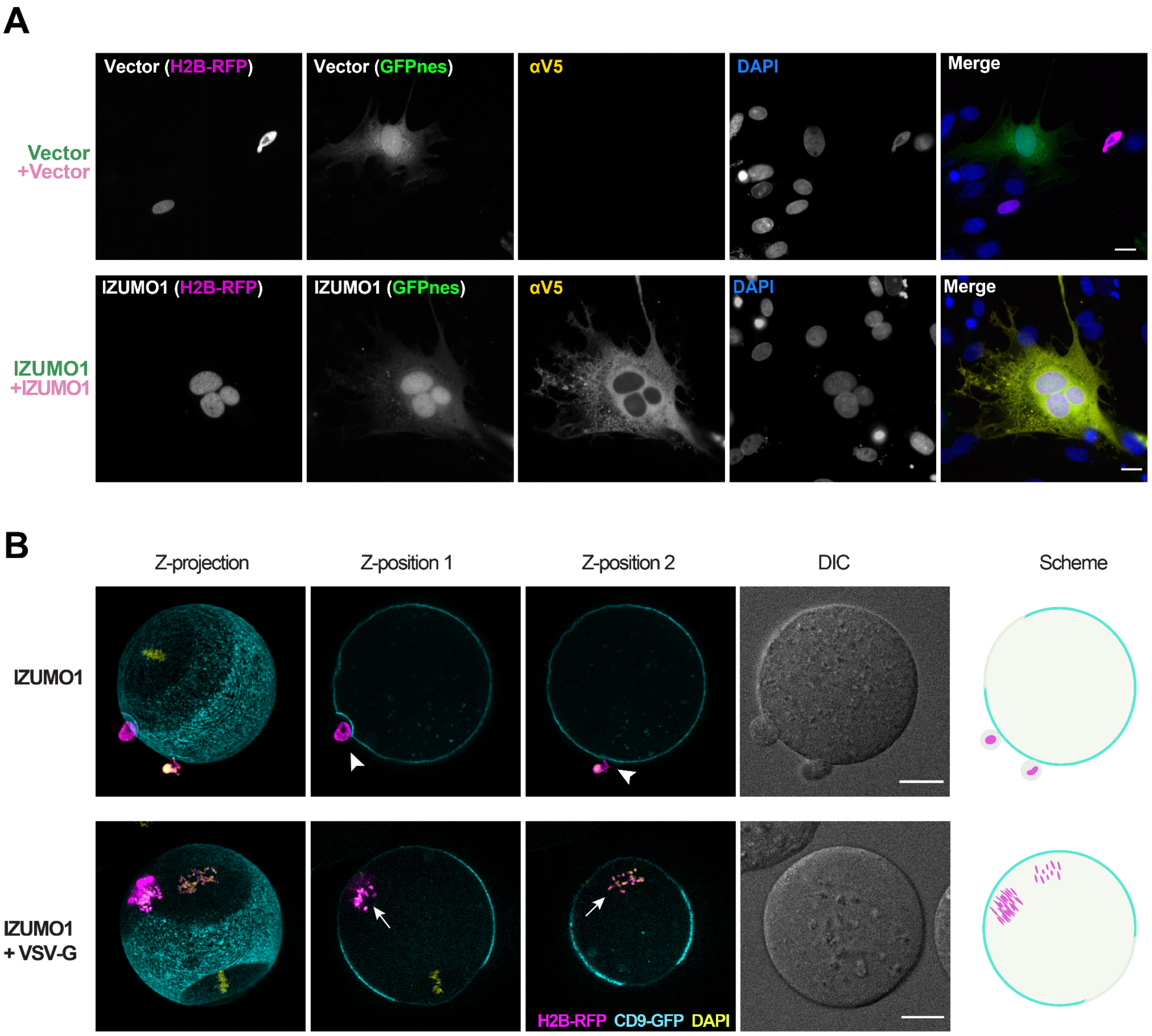


**Figure S4. Immunostaining of BHK cells after mixing experiments and BHK to oocyte fusion, related to Figures 2 and 5**

**(A)** Representative images of cells transfected with empty or IZUMO1-coding vectors after a content mixing assay and subjected to immunostaining using an anti-V5 antibody. Each separate channel (red, green, far red and DAPI) and merge are shown. Scale bars, 20 µm.

**(B)** BHK cells transfected with pCI::IZUMO1::H2B-RFP alone or together with a plasmid encoding for VSV-G were incubated with CD9-GFP oocytes (cyan) and observed by confocal microscopy. Fusion was evaluated as the presence of RFP-positive chromosomes (magenta) inside the cytoplasm of the eggs (arrows) as opposed to bound cells (arrowheads). For IZUMO1 no fusion was detected in >100 oocytes with cells attached, while for IZUMO1+VSV-G 18 out of 45 oocytes with cells bound displayed fusion. The maximal intensity Z-projection, two different focal planes (Z-position 1 and 2) and DIC are shown. DAPI staining is included in yellow. A schematic representation is shown on the right. Scale bars, 20 µm.

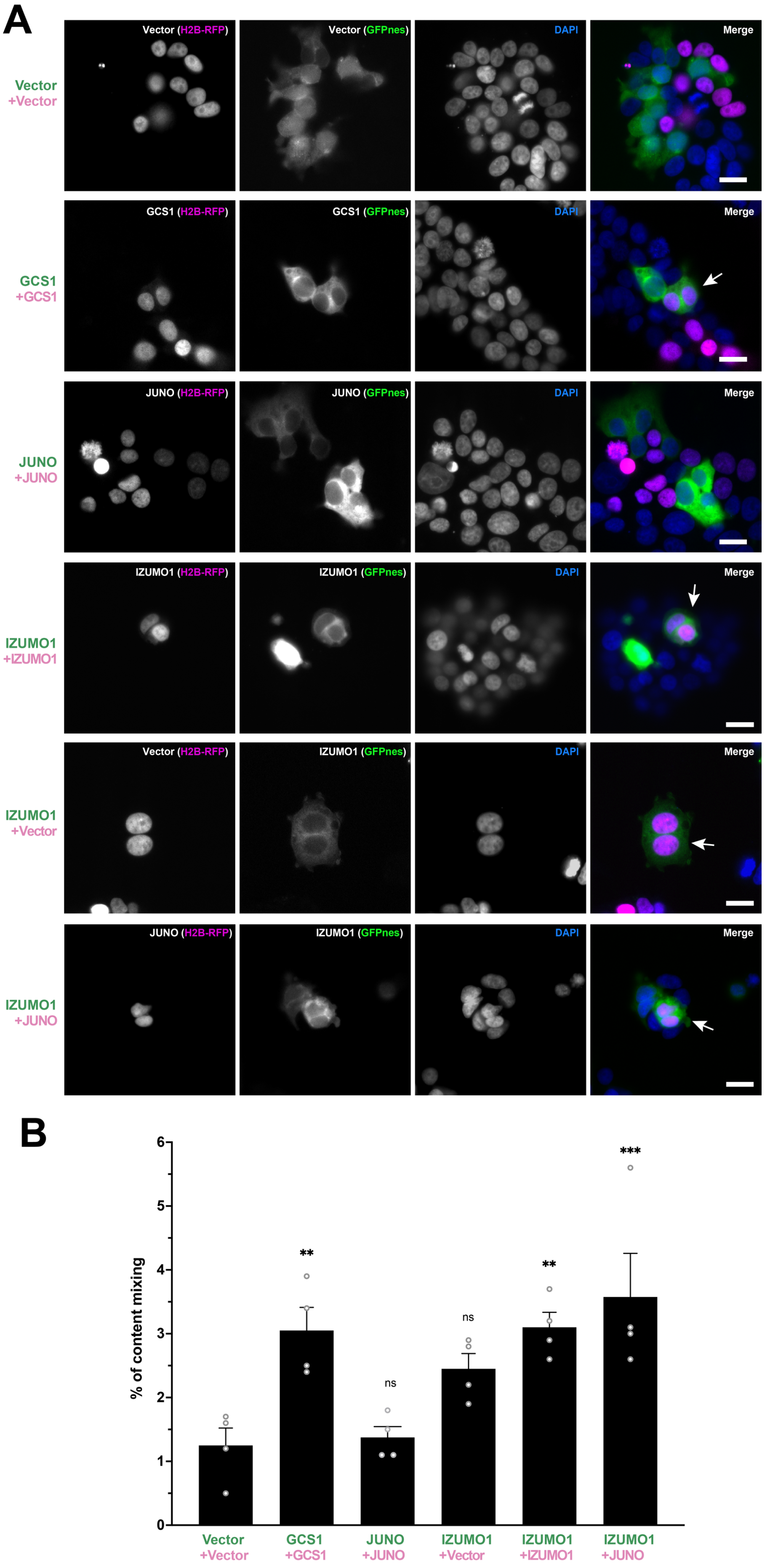
A
Vector (H2B-RFP)
Vector (GFPnes)
DAPI
Merge
Vector
+Vector
GCS1 (H2B-RFP)
GCS1 (GFPnes)
GCS1
+GCS1
JUNO (H2B-RFP)
JUNO (GFPnes)
JUNO
+JUNO
IZUMO1 (H2B-RFP)
IZUMO1 (GFPnes)
IZUMO1
+IZUMO1
Vector (H2B-RFP)
IZUMO1 (GFPnes)
IZUMO1
+Vector
JUNO (H2B-RFP)
IZUMO1 (GFPnes)
IZUMO1
+JUNO
B
% of content mixing
0
1
2
3
4
5
6
**
ns
ns
**
***
Vector +Vector
GCS1 +GCS1
JUNO +JUNO
IZUMO1 +Vector
IZUMO1 +IZUMO1
IZUMO1 +JUNO

**Figure S5. Evaluation of cell fusion by content mixing of HEK293T cells, related to Figure 2**

**(A)** Images of mixed cells transfected with pCI::GFPnes or pCI::H2B-RFP empty vectors or containing the coding sequence for the expression of GCS1/HAP2, IZUMO1 and JUNO as indicated in each panel. Arrows show fused cells containing both fluorescent markers, green cytoplasm (GFPnes) with red nuclei (H2B-RFP). DAPI staining is shown in blue. Each separate channel (red, green and DAPI) and merge are shown. Scale Bars, 20 μm.

**(B)** Quantification of content-mixing experiments with HEK293T cells. The percentage of mixing was defined as the ratio between the nuclei in mixed cells (NuM) and the total number of nuclei in mixed cells and fluorescent cells in contact that did not fuse (NuC), as follows: % of content mixing = (NuM/(NuM+NuC)) x 100. Bar chart showing individual experiment values and means ± SEM of four independent experiments. Comparisons by one-way ANOVA followed by Dunnett's test against the empty vectors. ns = non-significant, ** $p < 0.01$, *** $p < 0.001$.

### Movie legends

**Movie S1.** EFF-1-expressing cells show bilateral fusion corresponding to **Figure 3A**.
**Movies S2 and S3.** Two independent examples of IZUMO1-expressing cells showing bilateral fusion corresponding to **Figure 3A**.

### Supplementary Tables

**Table S1. GCS1/HAP2 and IZUMO1 induce syncytia formation.** Multinucleation was determined in four independent experiments in BHKs expressing myristoylated GFP (myrGFP), GCS1/HAP2, IZUMO1 or JUNO. Cells with 2 or more nuclei are considered multinucleated. Related to Figure 1.

| | Number of cells with: | | | | |
|---|---|---|---|---|---|
| | **1 nucleus** | **2 nuclei** | **3 nuclei** | **4 nuclei** | **5 nuclei** |
| **myrGFP** | 3462 | 180 | 8 | 0 | 0 |
| **GCS1/HAP2** | 2202 | 439 | 39 | 1 | 0 |
| **IZUMO1** | 2740 | 399 | 50 | 15 | 1 |
| **JUNO** | 3400 | 232 | 3 | 0 | 0 |

**Table S2. Primers used for this study**

| Primer name | Sequence 5'->3' | Description |
|---|---|---|
| ZP2 F EcoRI | GCGAATTCATGGCGAGGTGGCAGAGGAAAG | Forward primer for cloning *mZP2* into pIZT and pCAGGS with EcoRI |
| ZP2 R NotI | TGGGCGGCCGCCGTGATTGAACCTTATAGTTCTTTTC | Reverse primer for cloning *mZP2* into pIZT with NotI |
| ZP2 R NheI | CTTACGCTAGCTCAATGGTGATGGTGATGATG | Reverse primer for cloning *mZP2* into pCAGGS with NheI |
| Izumo1 F EcoRI | CCGGGAATTCGGTCGACTGGATCC | Forward primer for cloning *Izumo1* into pCAGGS with EcoRI |
| Izumo1 R XhoI | TCCATCTCGAGGGCCGCGTACG | Reverse primer for cloning *Izumo1* into pCAGGS with XhoI |
| Izumo1 F NheI | TTATCGCTAGCGAATTCGGTCGACTGGATCC | Forward primer for cloning *Izumo1* into pCI::H2B-RFP and pCI::GFPnes vectors with NheI |
| Izumo1 R SmaI | GTCCCCCGGGGCTCAATGGTGATGGTGATGATG | Reverse primer for cloning *Izumo1* into pCI::H2B-RFP and pCI::GFPnes vectors with SmaI |
| Juno F NheI | TAAGCTAGCCTCTTTGGCATCAGGAGGAGC | Forward primer for cloning *Juno* into pCI::H2B-RFP and pCI::GFPnes vectors with NheI |
| Juno R SmaI | ATACCCCGGGTGCCCCCAACATGAATAGCC | Reverse primer for cloning *Juno* into pCI::H2B-RFP and pCI::GFPnes vectors with SmaI |
| BlpI Flag F | TCAGCGACTACAAAGACGATGACGACAAGT | Forward oligo for tagging JUNO with a Flag tag using the BlpI site |
| BlpI Flag R | TGAACTTGTCGTCATCGTCTTTGTAGTCGC | Reverse oligo for tagging JUNO with a Flag tag using the BlpI site |
| GCS1 F NheI | CTAGCTAGCGGTACCATGGTGAACGCGATTTTAATG | Forward primer for cloning *GCS1/HAP2* into pCI::H2B-RFP and pCI::GFPnes vectors with NheI |
| GCS1 R SmaI | TCCCCCGGGCTAATGGTGATGGTGATGATGACC | Reverse primer for cloning *GCS1/HAP2* into pCI::H2B-RFP and pCI::GFPnes vectors with SmaI |
| GEX2 F KpnI | TCACAGGCCACCAAGCTTGGTACCATGGCGATTAAATT CGTTTCAC | Forward primer for generating and cloning *Gex2-venus* into pGENE vector with KpnI |
| GEX2 R | GCTGCCCCCTCCACCTGACTGCTTATTCTGGTTGCCGG AAG | Reverse primer for generating *Gex2-venus* |
| Venus F | TCAGGTGGAGGGGGCAGCGGGGGGGGAGGTATGGTG AGCAAGGGCGAG | Forward primer for generating *Gex2-venus* |
| Venus R NotI | GTGACCTCGAGCGGCCGCTTACTTGTACAGCTCGTCC ATGCC | Forward primer for generating and cloning *Gex2-venus* into pGENE vector with NotI |
| Izumo1-ecto-v5-R-SmaI | TCCCCCGGGCTACGTAGAATCGAGACCGAGGAGAGG GTTAGGGATAGGCTTACCTGGATTTTGAGCGACTGTA G | Reverse primer for cloning the ectodomain of Izumo1 with V5 tag and SmaI restriction enzyme site |
| Izumo1-W148A overlap F | ATGTCGCAGACTTTGATCGCTTGTCTTAAGTGCGAAAA G | Forward primer for cloning downstream of Izumo1$^{W148A}$ by overlap pcr |
| Izumo1-W148A overlap R | CTTTTCGCACTTAAGACAAGCGATCAAAGTCTGCGACA T | Reverse primer for cloning upstream of Izumo1$^{W148A}$ by overlap pcr |
| Izumo1-ΔIg overlap F | CGGAAATCCCTAGATTGTCCCCCAAAGCATTCAGAG | Forward primer for cloning downstream of Izumo1$^{\Delta Ig}$ by overlap pcr |
| Izumo1-ΔIg overlap R | CTCTGAATGCTTTGGGGGACAATCTAGGGATTTCCG | Reverse primer for cloning downstream of Izumo1$^{\Delta Ig}$ by overlap pcr |

| | | |
|---|---|---|
| Izumo1-F28A-F | TGCATCAAATGTGACCAGGCTGTGACAGATGCGCTAAAG | Forward primer for cloning the second fragment of Izumo1$^{FWW}$ mutant by overlap pcr |
| Izumo1-F28A-R | CTTTAGCGCATCTGTCACAGCCTGGTCACATTTGATGCA | Reverse primer for cloning the first fragment of Izumo1$^{FWW}$ mutant by overlap pcr |
| Izumo1-WWAA-R | CTTTTGATGACGAAGCATAGCCAATAGTTCCTTTATAAAGAGCTCTCCTTTTAAGTCACTGTCTGTAATACGCTTCAGATCCTTCAGAAAACTAGCGGTTGCTTGTTCCAGTGT | Reverse primer for cloning the second fragment of Izumo1$^{FWW}$ mutant by overlap pcr |
| Izumo1-W113A-F | TTTATAAAGGAACTATTGGCTATGCTTCGTCATCAAAAG | Forward primer for cloning the third fragment of Izumo1$^{FWW}$ mutant by overlap pcr |
| SmaI-His-R | TCCCCCGGGCTAATGGTGATGGTGATGATGACC | Reverse primer for cloning Izumo1 mutants |
| CD9-GFP F | TGAACCGCATCGAGCTGAAGGG | Forward primer for genotyping. For transgenic mice a 700 bp product is amplified. |
| CD9-GFP R | GAATATCACCAAGAGGAACC | Reverse primer for genotyping. For transgenic mice a 700 bp product is amplified. |
